# Local Inhibitory Dynamics Underpin Temporal Integration and Functional Segregation between Barrels and Septa in the Mouse Barrel Cortex

**DOI:** 10.1101/2024.01.23.576792

**Authors:** Ali Özgür Argunşah, Tevye Jason Stachniak, Jenq-Wei Yang, Linbi Cai, Alexander van der Bourg, Rahel Kastli, Theofanis Karayannis

**Affiliations:** Laboratory of Neural Circuit Assembly, Brain Research Institute, University of Zurich Winterthurerstrasse 190, CH-8057 Zurich, Switzerland; Neuroscience Center Zurich, University of Zurich and ETH Zurich, Winterthurerstrasse 190, CH-8057 Zurich, Switzerland; Division of Biomedical Sciences, Faculty of Medicine, Memorial University of Newfoundland, St. John’s, A1B 3V6, Canada; Department of Stem Cell & Regenerative Biology, Harvard University, Cambridge, MA, USA; University Research Priority Program (URPP), Adaptive Brain Circuits in Development and Learning, University of Zurich, Zurich, Switzerland

**Keywords:** Barrel Cortex, Elfn1 Knockout, Somatosensory Whisker Coding, Cortical Domains, Temporal Dynamics, Synaptic Integration, Sensory Projections, Progressive Functional Segregation

## Abstract

Just as separated digits and repeated sampling enhance somatosensation in humans, mice sense objects through multiple segregated whiskers through successive contacts. Individual whisker identity is maintained through the somatotopic organization of the Whisker→Brainstem→Thalamus→Cortex axis, culminating in distinct cortical domains: barrels and the surrounding septa. By performing simultaneous recordings using in-vivo electrophysiology in wild-type (WT) mice, we identify a progressive divergence in spiking activity between these domains upon repeated behaviorally relevant (10Hz) single- and multi-whisker stimulation. While the spiking activity ratio of multi- to single-whisker stimulation remains stable in barrels, it increases progressively in septa, suggesting inhibitory cell recruitment. Using genetic fate-mapping and tissue clearing, we indeed reveal that SST+ and VIP+ interneurons exhibit distinct laminar and regional distributions in barrel and septa domains. Further, calcium imaging of SST+ and VIP+ interneurons shows that while both neuron types respond to single-whisker stimulus, SST+ interneurons preferentially engage more in 10Hz multi-whisker stimulation, indicating their critical role in progressive stimulus preference. Genetic removal of Elfn1, which regulates the incoming excitatory synaptic dynamics onto SST+ interneurons, leads to the loss of the progressive increase in septal spiking ratio (MWS/SWS) upon stimulation. The importance of the loss of functional segregation of barrels, versus septa is revealed by cumulative temporal decoding analysis, supporting the notion that SST+ interneuron-mediated inhibition contributes to temporal encoding and stimulus integration. Finally, viral tracing combined with whole brain clearing and imaging reveals that barrel and septa domains project differentially to secondary somatosensory (S2) and motor (M1) cortices. These distinct projection patterns suggest that differential inhibitory processing in barrels and septa may contribute to functionally specialized downstream signaling. Together, our findings indicate that the progressive engagement of SST+ interneurons, mediated by Elfn1-dependent synaptic facilitation, underlies the preferential integration of multi-whisker stimuli in septa. This local inhibitory mechanism likely contributes to the functional segregation of barrel and septa domains and their distinct cortical projections, shaping how sensory information is processed and relayed to higher-order brain regions.

## Introduction

Although the somatosensory whisker system is one of the principal means by which mice sense their environment and navigate the world, these tasks are carried out by a relatively small number of whiskers. This small number of vibrissae is represented in the whisker somatosensory cortex as barrel “islands” (columns) separated and surrounded by the septal “sea” (compartments), reminiscent of the whisker pad pattern on a rodent’s face, where sparse hair follicles are separated by spaces between them. In contrast, other sensory systems, such as the auditory and the visual ones, have densely packed anatomical receptor configurations and cortical representations. Although all sensory cortices are composed of canonical microcircuits (Douglas and Martin, 2004) that include similar populations of excitatory and inhibitory neurons, maintaining segregated individual whisker information may require anatomical specializations, the barrel and septa domains. These domains may therefore create a distinct spatiotemporal stimulus representation and information coding in the barrel cortex, different from the other sensory modalities. In the adult murine barrel cortex, thalamocortical inputs coming from the ventro-postero-medial (VPM) and postero-medial (PoM) nuclei segregate into the barrel and septa domains, respectively (Alloway, 2008; Bureau et al., 2006; Kim and Ebner, 1999; Staiger and Petersen, 2021). Further, in mice (Audette et al., 2018; Sato and Svoboda, 2010; Shepherd and Svoboda, 2005) and rats (Alloway, 2008; Chakrabarti and Alloway, 2006; Melzer et al., 2006b) it was shown that information processing and flow within and from these domains target anatomically distinct sensory and motor areas. It has been suggested that the primary whisker somatosensory cortex (wS1) sends higher dimensional information needed for object recognition to the secondary somatosensory cortex (wS2), while the projections to the primary motor cortex (M1) carry less complex information that does not require segregated streams of information coming from individual whiskers (Alloway, 2008; Brecht and Sakmann, 2002; Cai et al., 2022; Sato and Svoboda, 2010). This divergence in information processing is also found within the barrel- and septa-related circuits. While barrel circuits are more involved in processing spatiotemporal whisker-object interactions, the septal circuits are more sensitive to the frequency of whisker movements, which suggests the use of temporal versus rate coding of information in barrel and septa, respectively (Melzer et al., 2006a, 2006b). Hence the existence of the septal column separating the barrels may provide mice with the means to separate these two interconnected yet very distinct information-processing routes along the temporal domain, and to make sense of their somatosensory environment through repeated sampling via cloistered information streams. Recent studies in mice have shown that barrel and septa compartments are not only anatomically segregated in terms of cell-type specific thalamocortical projections and differential expression of different genetic markers (Young et al., 2023) but also in their tuning upon single- and multi- whisker stimulation (Wang et al. 2022).

In both rats and mice, the operational identity of barrels and septa has often been attributed to the parallel bottom-up thalamocortical pathways of VPM and PoM. Here, by performing *in-vivo* silicon probe recordings simultaneously in the barrel and septa domains upon repeated single- and multi- whisker stimulation in wS1 of mice, we first show and characterize the temporal divergence between these domains. Utilizing genetic fate-mapping, passive tissue clearing, and light-sheet microscopy, we reveal that cortical somatostatin (SST+) and vasoactive intestinal peptide (VIP+) expressing interneurons show a layer-dependent differential distribution in the barrel versus septa columns. Through two-photon calcium imaging of these two populations, we show a differential engagement of SST+ cells upon multi- vs. single-whisker stimulation. By altering the short-term synaptic dynamics of incoming excitation onto these interneurons through the removal of the trans-synaptic protein Elfn1 (Elfn1 KO), we find that the domain-specific contrast ratio between multi- over single- whisker response to a repetitive stimulus is diminished. Through a progressive decoding analysis, we find that in contrast to control, whisker stimulation-evoked Elfn1 KO responses in different domains cannot be decoded efficiently. Finally, using *in-vivo* retrograde viral-based tracing, we show a layer-specific projection preference between these domains and downstream regions wS2 and M1. Hence, in addition to the anatomical specialization through thalamocortical pathways, we reveal a key contribution of local lateral cortical inhibition, provided by SST+ interneurons, in setting up the functional segregation of barrel and septa domains, and subsequently, their downstream targets.

## Results

### Barrel and Septa columns display differentially adaptive spiking upon repeated single- and multi- whisker stimulation

To characterize the temporal stimulus-response profiles of barrel and septal domains simultaneously, we performed *in-vivo* multi-shank (8x8) silicon probe recordings under urethane-induced light anesthesia from mouse somatosensory whisker barrel cortex (wS1), after the age of active whisking onset, postnatal day (P)20-30. Whisker stimulation was performed on either just one principal whisker (single-whisker stimulus or SWS) or via both the principal whisker and most of the macro vibrissae together (multi-whisker stimulus or MWS; see methods) (Figure 1A). Single- or multi- whisker evoked activity within identified regions was recorded upon a 2s-long stimulation at 10Hz, similar to the average frequency at which mice actively whisk (Carvell and Simons, 1990; Grant et al., 2012). Each mouse received both stimulation paradigms. Current source density (CSD) analysis was used to assess the location of each silicon probe shank, which was later verified by post-hoc histology. The recorded activity was assigned to the stimulated principal barrel column (B), the adjacent septal column (S), or the adjacent unstimulated neighboring barrel column (N) according to SWS paradigm (Figure 1B and Supp. Fig. 1A to D). An example probe assignment can be seen in Supp. Fig. 1E. While this distinctive experimental paradigm is relatively low throughput, it has allowed us to investigate all three of these domains simultaneously. The septal compartments in mouse barrel cortex are quite narrow (Bureau et al., 2006; Sato et al., 2007) and there is the possibility of rotational misalignment during the insertion of silicon probes into the brain (Supp. Fig. 1F). For these reasons we restricted our analysis of *in-vivo* responses upon SWS and MWS between barrel and septa responses to upper layers and L4, for which we have higher confidence about the anatomical domain allocation after our histological (Supp. Fig. 1B) and CSD analysis (Supp. Fig. 1C and D). In line with this domain allocation, t-distributed stochastic neighbor embedding (t-SNE) analysis of the multi-unit activity (MUA) recorded across the entire train of stimuli in the different domains revealed that the spiking responses between the principal barrel and the neighboring one form two separate clusters, while the responses recorded in the septa lay in between (Figure 1C; see methods). Accordingly, we interpret multi-unit activity recorded from septal electrode positions as reflecting septal-enriched populations rather than exclusively septal neurons, while emphasizing that the conclusions are based on relative differences between domains under matched recording conditions.

**Figure 1.**
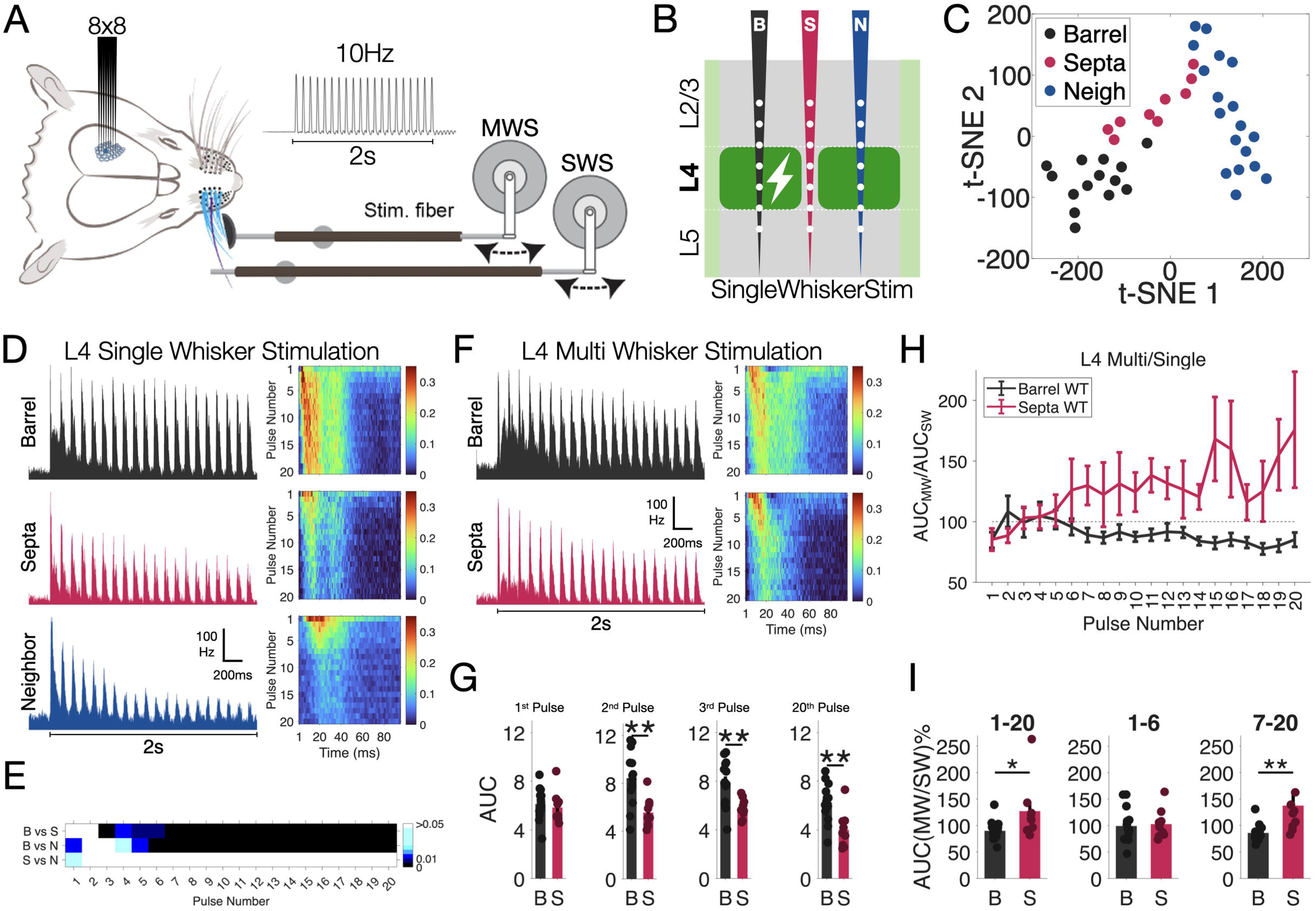
Temporal divergences emerge between barrel and septa domains upon repeated whisker stimulation in Layer 4 of mouse wS1. (A) Either the principal whisker (SWS, or single whisker) or most of the whiskers (MWS, or multi whisker) were stimulated for 2s at 10Hz (20 pulses) for 20 times (20 trials). (B) Acute simultaneous *in-vivo* silicon probe (8x8) recordings were performed from barrel and septa domains (>P21) (C) t-SNE analysis of the average firing dynamics from three domains after averaging 20 single trials. (D) Left: Average multi-unit firing profiles recorded at Layer 4 of barrel (black), unstimulated neighbor (blue) and septa (pink) columns upon 2s-long 10Hz repeated single whisker stimulation. Right: Pulse-aligned firing heatmap of the SWS data. Color scale bar represents spikes per millisecond. (E) Pair-wise statistical comparisons of first 50ms of the single whisker responses from barrel (B), septa (S) and neighbor (N) for each pulse (Mann-Whitney-U-Test). (F) Left: Average multi-unit firing profiles recorded at Layer 4 of barrel (black) and septa (pink) columns upon 2s-long 10Hz repeated multi whisker stimulation. Right: Pulse-aligned firing heatmap of the MWS data. Color scale bar represents spikes per millisecond. (G) Pair-wise statistical comparisons of first 50ms of the multi whisker responses from barrel (B) and septa (S) for the 1^st^, 2^nd^,3^rd^ and 20^th^ pulses (Mann-Whitney-U-Test). (H) Temporal ratio dynamics (average AUC MW responses divided by average AUC SW responses) for barrel and septa. (I) statistical comparison of average AUC ratios of all pulses (1-20), average of the first six (1-6) or average of the last 14 pulses (7-20), respectively. (p values are color coded in E, stars represent *: p<0.05; **: p<0.01; ***: p<0.001 in G and I.)

It has previously been shown that electrophysiological responses upon whisking-based object detection saturate rapidly after a single whisker touch, while object localization may require multiple repeated whisker touches (O’Connor et al., 2013; Pammer et al., 2013). Correspondingly, when a single whisker is stimulated repeatedly, the response to the first pulse is principally bottom-up thalamic-driven responses, while the later pulses in the train are expected to also gradually engage cortico-thalamo-cortical and cortico-cortical loops (Kyriazi and Simons, 1993; Middleton et al., 2010; Russo et al., 2025). We therefore first tested whether the leading pulses of the stimulus train evoked differential responses among the principal barrel, the adjacent septa, and the neighboring barrel upon SWS and MWS. For easier visualization and full appreciation of the data, we present our findings both in classical time series firing-rate plot and stimulation pulse aligned stacked images (Supp. Fig. 2A and B, 200ms baseline followed by 2000ms evoked response to 20 pulses at 10Hz reshaped to 20 stacked 100ms-long evoked response image, left to right). Earlier studies that separately recorded whisker-evoked responses from either barrel or septa domains in rat reported that the responses of barrel neurons to single principal whisker stimulation are much higher than those evoked in the septal neurons (Armstrong-James and Fox, 1987; Brecht and Sakmann, 2002). The analysis of the data we obtained from the simultaneous recordings at these domains in the mouse do not reveal such differences in the first two stimulus responses in L4 upon single whisker stimulation (SWS) (Figure 1D and 1E; white represents no significance). However, response differences emerged from the 3^rd^ pulse onward upon SWS (Figure 1E). Similar differences also emerged between stimulated principal and unstimulated neighboring barrel columns upon SWS (starting from 4^th^ pulse, Figure1E, Barrel-vs-Neighbor (BN) row). Interestingly, an inverted pattern of significance appeared between septa and adjacent unstimulated neighboring barrels (Figure 1E, Septa-vs-Neighbor (SN) row). Additionally, repeated-measures ANOVA (rmANOVA) analysis revealed that a significant main effect of Condition (F(2, 38) = 18.322, p < 0.001) and a significant Condition-Time interaction (F(38, 722) = 14.618, p < 0.001), indicating robust differences in overall responses and their temporal dynamics across the conditions upon SWS. When multiple whiskers (MWS) were activated simultaneously, like SWS, a divergence between barrel and septa domain activity also occurred in Layer 4 from the 2^nd^ pulse onward (Figure 1F and 1G). However, while, rmANOVA revealed a significant main effect of Condition (F(1, 21) = 25.666, p < 0.001), it showed a non-significant Condition-Time interaction (F(19, 399) = 1.9076, p = 0.13229), indicating robust overall differences in activity between the conditions, but no significant variation in their temporal patterns across the 20 time points. Finally, when we calculated the response ratio upon MW over SW stimulation, we saw that the septal MW/SW ratio diverged from barrel MW/SW ratio over the course of repeated stimulation (Figure 1H and 1I), indicating a role of septal neurons in the progressive processing MW information. This result was later confirmed by rmANOVA, which revealed a significant main effect of Condition (F(1, 21) = 5.6399, p = 0.027162) and a significant Condition-Time interaction (F(19, 399) = 4.9023, p = 0.011313), supporting differences in overall responses and their temporal dynamics between the conditions. Post-hoc tests confirmed significant differences between the multi/single rations of Barrel and Septa data at multiple time points (e.g., p < 0.0025 at times 3, 4, 6, 7, 8, 10, 11, 12, 16, 19 after Bonferroni posthoc correction). The persistence of domain-specific effects in Layer 4 at a higher spike-detection threshold (S.D.>9.5) supports the interpretation that these responses arise from neuronal populations spatially close to the recording sites, rather than from broad mixing across adjacent domains (Supp. Fig. 3).

### Barrel and Septa columns display differential SST+ and VIP+ neuron densities

It is known that barrel and septa domains receive different thalamic projections, from VPM and PoM respectively, that might affect stimulus response properties (Ahissar et al., 2001; Chmielowska et al., 1989; Koralek et al., 1988; Sosnik et al., 2001). Our electrophysiological experiments show a significant divergence of responses between domains upon both SWS and MWS in L4. We hypothesized that this late divergence might be driven by differences in the local circuitry, and specifically inhibitory cells. We therefore next assessed the spatial distribution of two inhibitory cell populations that could provide distinct regulation of cortical activity in the temporal domain: SST+ and VIP+ cells. Earlier research examining the three largest inhibitory neuron populations in the mouse barrel cortex showed that, while parvalbumin positive (PV+) cells do not show any differential density preference between barrel and septa columns, SST+ and VIP+ cells have higher density in the septa in L4 (Almási et al., 2019). We attempted to verify the reported differential SST+ and VIP+ neuronal distributions using an alternative imaging and counting strategy. To label the two neuronal populations, we crossed the SST-Cre and VIP-Cre lines with a tdTomato reporter mouse line (Ai14). To comprehensively quantify the density of SST+ and VIP+ cells in the barrel and septa domains in 3D, we utilized a passive CLARITY-based tissue clearing protocol, followed by light-sheet microscopy using a custom-built light-sheet microscope (Vladimirov et al., 2023; Voigt et al., 2019) (Figure 2A). The individual barrels in wS1 were reliably detected using auto-fluorescence from the tissue acquired with a 488nm laser. tdTomato positive SST+ or VIP+ neurons were detected by a 561nm laser, allowing us to accurately localize and count cells in barrel and septal domains (Figure 2B and C). By adjusting the angle of each barrel column in 3D (using Imaris, RRID:SCR_007370), we precisely identified the column, using the barrel borders for every plane (XY, YZ, XZ) in L4 (Figure 2B, also see Methods). Given that the depth of L4 can be reliably measured due to its well-defined barrel boundaries, and that the relative widths of other layers have been previously characterized (Lefort et al., 2009), we estimated laminar boundaries proportionally. Specifically, Layer 2/3 was set to approximately 1.3–1.5 times the width of L4, Layer 5a to ∼0.5 times, and Layer 5b to a similar width as L4. Assuming isotropic tissue expansion across the cortical column (Ueda et al., 2020), we extrapolated the remaining laminar thicknesses proportionally. Once the columns were accurately determined, we counted the number of tdTomato positive neurons in the barrels and septa and corresponding volumes for density estimation. Despite their opposite distributions and neuronal density changes along the depth of wS1 (Figure 2B and C), we found that both SST+ (Figure 2D) and VIP+ (Figure 2E) neuronal densities were higher in L4 septa compared to barrels, but we detected no difference in L2/3 and L5.

**Figure 2.**
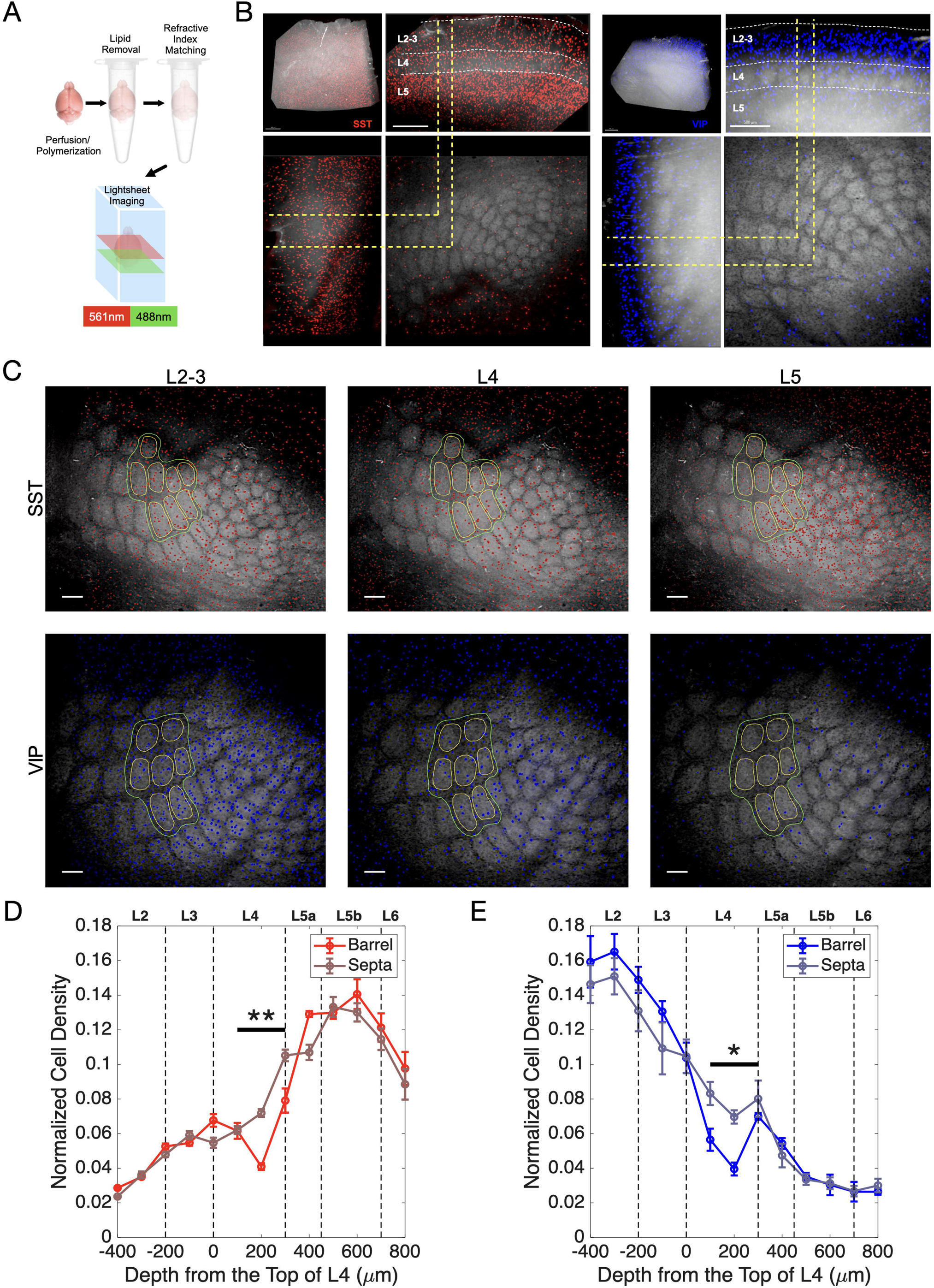
SST+ and VIP+ neuron densities differ at Barrel and Septa Domains in Layer 4. (A) A passive CLARITY-based tissue clearing protocol was performed on the collected brains which were subsequently imaged in their entirety using a custom-built light-sheet microscope (mesoSPIM). 561nm wavelength was used to image tdTomato, 488nm wavelength was used to image autofluorescence. (B) 3D reconstruction of cleared SST-Ai14 (left) and VIP-Ai14 (right) barrel cortex. X-Y, X-Z and Y-Z views are presented focusing with a single barrel max projection. The brain was rotated to achieve the orientation that the barrel columns are perpendicular to the X-Y view. The scale bars are set for 500μm. (C) Max projection examples of SST (red) and VIP (blue) neurons in wS1 L2-3, L4 and L5. Quantified barrels (yellow lines) and septa (green lines) are circled out which are perpendicular to X-Y (top-down) view. The scale bar is set for 200μm. (D) SST+ neuron density distribution over laminae for barrel and septal domains (N=4 mice, n=54 Barrels and surrounding septa per barrel). Normalized cell densities plotted as a function of depth. 0µm represents the top of layer 4. SST+ interneuron density is significantly higher in septa (p=0.0015, F=30.64, repeated-measures ANOVA). (E) VIP+ neuron density distribution over laminae for barrel and septal domains. (N=4 mice, n=53 Barrels and surrounding septa per barrel). Normalized cell densities plotted as a function of depth. 0µm represents the top of layer 4. VIP+ interneuron density is also significantly higher in septa (p=0.021, F=9.63, repeated-measures ANOVA).

These results confirmed that SST+ and VIP+ interneurons have higher densities in septa compared to barrels in L4 and suggest these interneurons may play a role in diverging responses between barrels and septa.

### SST+ neurons are more strongly activated by multi- versus single-whisker stimulation

Having identified spatial distribution patterns of SST+ or VIP+ interneurons that could contribute to the single- vs multi- whisker response scaling observed in Figure 1F; we sought to assess these interneurons’ response profiles upon similar whisker stimulation paradigms. Therefore, we performed similar SWS and MWS experiments using acute in-vivo two-photon calcium (Ca2+) imaging under light anesthesia after weaning (P21–41) (Figure 3A and B). We injected OGB-1, the membrane-permeable AM-ester form of the calcium indicator, into wS1 of animals expressing tdTomato in either VIP+ or SST+ INs (VIPCre-Ai14 and SSTCre-Ai14 lines, respectively) to localize VIP+ and SST+ activity (Figure 3A). Two-photon Ca2+ imaging was performed upon stimulation of either the C2 whisker at 10Hz for 2s (SWS) or most of the macro vibrissae including C2 (MWS) (Figure 3B). We found that both SST+ and VIP+ interneurons increased their firing upon SWS (Figure 3C and D) as has been reported previously (Kastli et al., 2020), but only SST+ interneurons changed their firing significantly between single- and multi- whisker stimulation at 10-Hz (Figure 3D, SW vs MW comparison). Although these optical recordings were performed in layer 2/3 (L2/3) rather than layer 4 (L4) due to depth limitations, they provide independent, single-cell–level evidence that SST⁺ interneurons are preferentially recruited by multi-whisker stimulation in a stimulus-dependent manner. This supports a cell-type–specific role for SST⁺ interneurons in shaping temporal integration during repeated sensory stimulation, complementing the Layer-4 population effects revealed by electrophysiology, while not implying spatially focal domain specificity at the level of L2/3 multi-unit signals.

**Figure 3.**
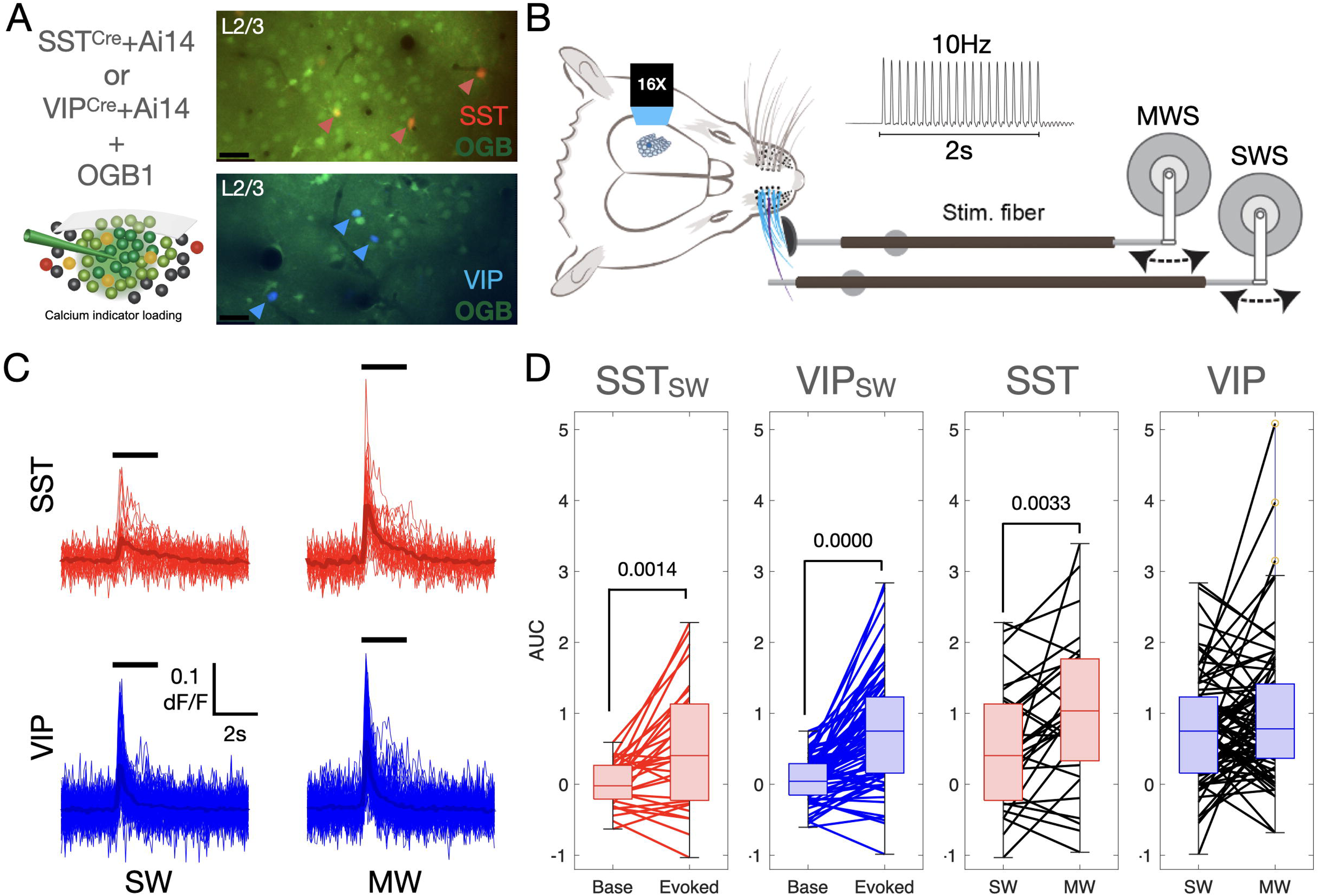
Single- vs multi- whisker responses of SST+ and VIP+ Interneurons (A) Interneurons are labeled with tdTomato using reporter mouse lines and Ca^2+^ imaging has been performed after bulk loading of OGB-1. scale bar 35μm. (B) Schematic representation of the single- and multi- whisker stimulation protocol upon 10Hz stimulation. (C) Δ*F/F* traces of evoked activity of SST (on top) and VIP (on bottom) interneurons (*N* = 3 animals per group, VIP P21+: 138 cells, SST P21+: 51 cells). (D) Area under the curve (AUC) of the first 2s of the Δ*F/F* traces of all SST and VIP cells upon single- (SW: 2s-long principal whisker evoked activity) and multi- whisker (MW: 2s-long multi whisker stimulation evoked activity) stimulation (Wilcoxon signed-rank test, Baseline: 2s-long baseline activity).

### Progressive multi- over single-whisker response divergence in septal populations is abolished in Elfn1 KO mice

Building on prior evidence that Elfn1 knockout disrupts short-term facilitation in SST+ interneurons (Sylwestrak and Ghosh, 2012; Tomioka et al., 2014; Stachniak et al., 2019, 2021, 2023), we attribute the abolished barrel-septa divergence in Elfn1 KO mice to altered SST+ synaptic dynamics, though direct synaptic measurements were not performed here. SST+ interneurons in the cortex are known to show distinct short-term synaptic plasticity, particularly strong facilitation of excitatory inputs, which enables them to regulate the temporal dynamics of cortical circuits (Grier et al., 2023; Liguz-Lecznar et al., 2016). This facilitation allows SST+ cells to progressively enhance their inhibitory influence in response to repeated stimulation, a property critical for shaping network activity. A key regulator of this plasticity is the synaptic protein Elfn1, which mediates short-term synaptic facilitation of excitation on SST+ interneurons (Sylwestrak and Ghosh, 2012; Tomioka et al., 2014).

Having identified SST+ cells as potential regulators of temporal MW/SW ratio divergence, we investigated how the absence of Elfn1 alters sensory-driven activity in vivo, potentially shaping septal function in MW/SW response scaling. Using the same whisker stimulation paradigms and domain-assignment strategy as in WT animals (Figure 1A and B; and Supp. Figure 1), we recorded from barrel and septa domains in Elfn1 KO mice under both single-whisker stimulation (SWS) and multi-whisker stimulation (MWS) conditions. Unsupervised t-SNE analysis revealed that the domain-specific clustering of responses seen in WT (Figure 1C) mice was disrupted in Elfn1 KO mice, with barrel and septa responses intermingling (Figure 4A). This convergence was also evident in stimulus-aligned stacked images, where the distinct differences between barrels and septa under SWS were largely abolished in the KO (Figure 4B, C). This was reflected in the ANOVA results, which showed a non-significant main effect of Condition (F(2, 26) = 1.76, p = 0.192) but a significant Condition-Time interaction (F(38, 494) = 10.26, p < 0.001), indicating that although the overall activity levels were similar, the temporal dynamics differed across conditions. The only remaining distinction in the KO was between directly stimulated barrels and neighboring barrels, which persisted (Figure 4B, C, Barrel-vs-Neighbor row). Interestingly, although paired analysis showed that the temporal divergence between barrel and septal domains observed in WT Layer 4 was diminished in Elfn1 KO mice (Figure 4D, E), rmANOVA revealed that both the main effect of Condition (F(1, 16) = 13.39, p = 0.0021) and the Condition-Time interaction (F(19, 304) = 6.50, p = 0.00034) were significant—unlike the corresponding MWS data in wild-type animals. However, the progressive increase in the MW/SW response ratio seen in septal neurons of WT mice was absent in KO mice (Figure 4F, G, paired test), with rmANOVA showing a non-significant main effect of Condition (F(1, 16) = 0.22, p = 0.647) and a non-significant Condition × Time interaction (F(19, 304) = 1.17, p = 0.281). These results together indicate that the barrel–septum difference observed in WT animals was lost in the Elfn1 KO.

**Figure 4.**
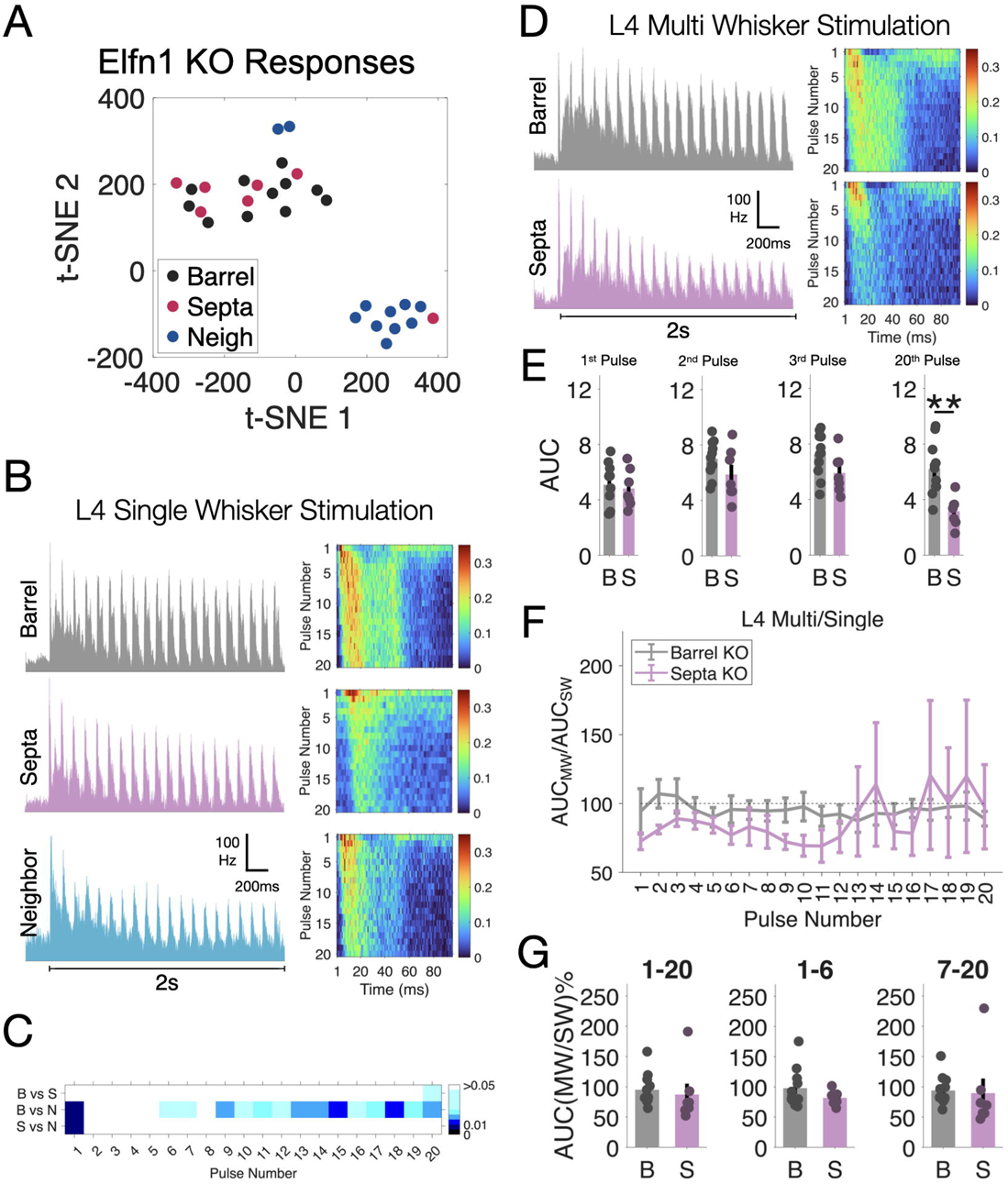
Loss of Elfn1 abolishes barrel-septa response divergences upon single-whisker stimulation. (A) t-SNE analysis of the average firing dynamics from three domains. (B) Left: Average multi-unit firing profiles recorded at Layer 4 of barrel (gray), unstimulated neighbor (light blue) and septa (purple) columns (11-barrel columns, 7 septal columns, 11 unstimulated neighbors from N=4 mice) upon 2s-long 10Hz repeated single whisker stimulation. Right: Pulse-aligned firing heatmap of the same data by representing each stimulation pulse response on one row. Color scale bar represents spikes per millisecond. (C) Pair-wise statistical comparisons of first 50ms of the single whisker responses from barrel (B), septa (S) and neighbor (N) for each pulse (Mann-Whitney-U-Test). (D) Left: Average KO multi-unit firing profiles recorded at Layer 4 of barrel (gray) and septa (magenta) columns upon 2s-long 10Hz repeated multi whisker stimulation. Right: Pulse-aligned firing heatmap of the KO MWS data. Color scale bar represents spikes per millisecond. (E) Pair-wise statistical comparisons of first 50ms of the multi whisker responses from barrel (B) and septa (S) for the 1^st^, 2^nd^,3^rd^ and 20^th^ pulses (Mann-Whitney-U-Test). (F) Temporal ratio dynamics (average AUC KO MW responses divided by average AUC KO SW responses) for barrel and septa. (G) statistical comparison of average AUC ratios of all pulses (1-20), average of the first six (1-6) or average of the last 14 pulses (7-20), respectively. (p values are color coded in C, stars represent **: p<0.01 in E.)

These findings suggest that in WT animals, activity spillover from principal barrels is normally constrained by the progressive engagement of SST+ interneurons, driven by Elfn1-dependent facilitation at their excitatory synapses. In the absence of Elfn1, this local inhibitory mechanism is disrupted, leading to a loss of the distinct temporal response divergence between barrel and septa domains.

### Barrel and Septa response decoding identity is lost in Elfn1 KO mice

Based on our observation that Elfn1 is essential for sustaining barrel-septa response divergence, we sought to quantify how sensory information accumulates and is integrated over time in these domains—and how this process is altered in Elfn1 KO mice.

To investigate this, we performed an accumulative temporal decoding analysis using neural responses to 20 pulses of either single-whisker stimulation (SWS) or multi-whisker stimulation (MWS) at 10 Hz for 2 seconds. We segmented the response window into three time ranges to identify the temporal segments driving divergence between domains: the full response window (1-95ms), the first part (1-50ms) which includes the first peak, and the late part (51-95ms), which includes the second firing peak (Figure 5A). A one-versus-all error-correcting output code (ECOC) classifier, using a GentleBoost ensemble of decision trees (Allwein et al., 2001; Friedman et al., 2000), was trained on firing profiles from WT barrel, septa, and neighboring barrel domains (3-class classification for SWS) or barrel and septa domains (2-class classification for MWS). We conducted 10-fold cross-validation within WT and KO datasets (WT CrossVal and KO CrossVal) and tested a WT-trained classifier on Elfn1 KO responses (WT Train - KO Test) to evaluate decoding performance across accumulated pulses (see Methods).

**Figure 5.**
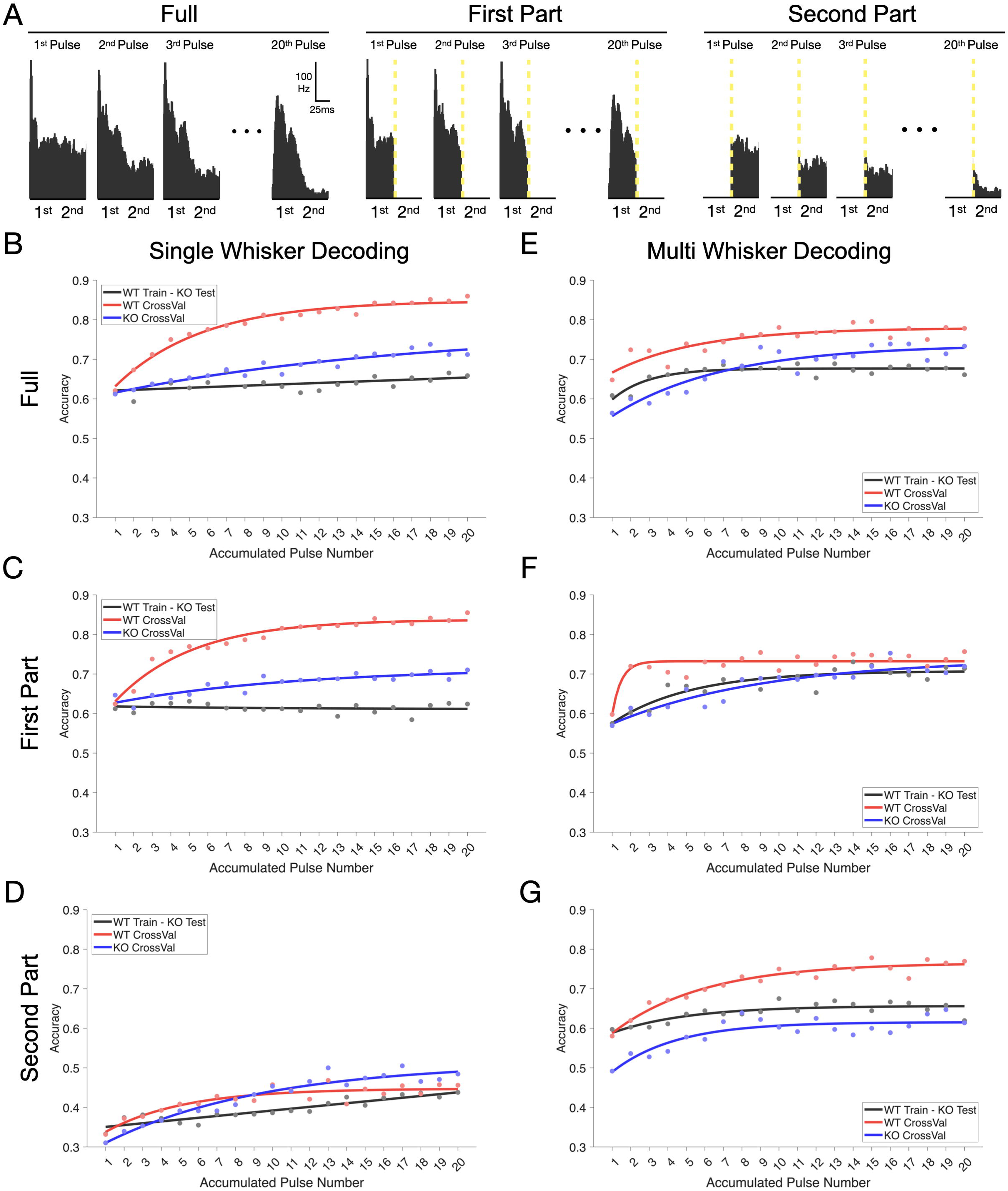
Accumulative decoding analysis shows an alteration of columnar domain identity in Elfn1 KO animals. (A) For the decoding analysis, the entire pulse response, only the first 50ms or only the last 45ms has been used. (B) Decoder analysis of the full pulse response profiles in L4 upon SWS. (C) Decoder analysis of the first part of the pulse response (1-50ms) profiles in L4 upon SWS. (D) Decoder analysis of the second part of the pulse response (51-95ms) profiles in L4 upon SWS. (E) Decoder analysis of the full pulse response profiles in L4 upon MWS. (F) Decoder analysis of the first part of the pulse response (1-50ms) profiles in L4 upon MWS (G) Decoder analysis of the second part of the pulse response (51-95ms) profiles in L4 upon MWS.

In WT mice, decoding accuracy increased progressively with the number of pulses upon SWS (Figure 5B, red) reflecting accumulated segregation of sensory information over time. This SWS WT CrossVal trend was more pronounced in the first response window (1-50ms) (Figure 5C vs 5D, red), indicating that domain-specific information accumulates predominantly in first temporal segments in repeated single whisker sampling. Cross-validation within the WT dataset (WT CrossVal) showed higher accuracy compared to the WT-trained classifier tested on KO responses (WT Train - KO Test) and KO CrossVal at this first segment, but not in the second (Figure 5B-D). Elfn1 KO dataset showed higher cross validation performance at the second part of the response profiles (Figure 5D) indicating a temporal information delay under the reduced progressive inhibition. While decoding curves started at similar initial levels for SWS data (Figures 5B-D, first pulse number), initial MW decoding accuracies were remarkably different (Figures 5E-G). In WT mice, while the decoding accuracy increased progressively as in SWS (Figure 5E, red), the difference between accumulated 20^th^ and the starting (1^st^) accuracies were lower than KO CrossVal results (Figures 5E-G, red vs blue) which was predominantly due to the first part of the signal (Figure 5F vs 5G).

To assess whether stimulus representations generalize between single-whisker stimulation (SWS) and multi-whisker stimulation (MWS), we trained ECOC classifiers (GentleBoost) on MWS responses and tested them on SWS data, evaluating decoding accuracy for barrel and septal compartments in layer 4 (L4) of the barrel cortex (Supp. Fig. 5A–C). Over the full response window (1–95ms; Supp. Fig. 5A), both WT and KO animals showed high generalization, with septal accuracy reaching 1.0 by pulse 6 in WT and pulse 8 in KO. Septal accuracy was higher in WT (e.g., 0.8167 vs. 0.7679 for barrels at pulse 1), while in KO, barrels initially led (0.8000 vs. 0.7571 at pulse 1) before septa surpassed by pulse 4 (0.9214 vs. 0.9182).In the early epoch (1–50ms; Supp. Fig. 5B), WT septal accuracy was slightly higher (0.7556 vs. 0.7143 for barrels at pulse 1), converging to 1.0 by pulse 8. In KO, barrel accuracy exceeded septa at pulse 1 (0.8045 vs. 0.7500), but septa reached 1.0 by pulse 10, while barrels plateaued at 0.9955. In the later epoch (51–95ms; Supp. Fig. 5C), septal accuracy surpassed barrels in both genotypes (WT: 0.9944 vs. 0.9214; KO: 0.9714 vs. 0.9227 at pulse 20), with the advantage evident from pulse 1 (WT: 0.7389 vs. 0.6286; KO: 0.6357 vs. 0.6364) and growing with pulses.

These findings indicate that septa process SWS and MWS differently, with higher decoding accuracy reflecting distinct, posterior medial nucleus (POm)-driven MWS responses compared to minimal SWS activation. Barrels, driven by consistent ventral posteromedial nucleus (VPM) input, show less distinctiveness, yielding lower accuracy. In Elfn1 KO mice, the initial barrel advantage (1–50ms) suggests reduced early septal distinctiveness due to disrupted excitatory drive to somatostatin-positive (SST+) interneurons, with recovery in later pulses indicating compensatory mechanisms. Calcium imaging confirms stronger SST+ interneuron activation in septa during MWS, driving late-phase (51–95ms) response differences amplified by successive pulses. These results highlight Elfn1’s role in temporal integration, where SST+ interneurons enhance septal MWS processing, maintaining functional segregation between domains.

### wS1 Barrel Columns and Septal Domains have different wS2 and M1 Layer-dependent projection preferences

We identified that temporal MW/SW response divergence in septal populations is abolished in Elfn1 KO mice. Given that barrel and septal compartments differentially process sensory input over time, this raises the question of how these distinct streams of information are relayed to downstream regions and whether they contribute to separate functional pathways. Following up from this observation, we finally aimed to explore if the projection targets of these two wS1 domains are distinct, in a similar manner to what has been revealed for visual cortex domain projections (Meier et al., 2021). Previously, it has been shown that cortical pyramidal neurons in wS1 exhibit distinct long-range projection patterns and sensory tuning properties depending on their projection target (S2 or M1). S2-projecting pyramidal neurons tend to have narrower receptive fields, are more likely to be tuned to the columnar whisker and carry more precise sensory information about whisker deflections. In contrast, M1-projecting pyramidal neurons are more broadly tuned, integrating information from multiple whiskers, which aligns with their role in sensorimotor integration and behavior (Sato and Svoboda, 2010; Yamashita et al., 2013). This data could be suggestive of distinct localization of the different projection neurons in the different wS1 domains. By utilizing *in vivo* retrograde AAV injections in two of the main target cortical regions of wS1, we aimed to characterize the projections from wS1 to wS2 and M1.

We injected a retrograde AAV virus expressing GFP in wS2 or M1 in separate adult mice and dissected the brains 4 weeks after the injection (Figure 6A). We utilized again the passive CLARITY-based tissue clearing and imaged the whole brain using mesoSPIM. The auto-fluorescence of the barrels allowed for labeling of barrel columns in 3D (Figure 6B). With the same strategy as our previous analysis on interneuron population distributions (Figure 2), we counted the number of neurons in wS1 barrel and septa domains across the cortical layers (Figure 6C) and measured the corresponding domain volumes to estimate projection densities as a proxy for connectivity levels between S1-S2 and S1-M1.

**Figure 6.**
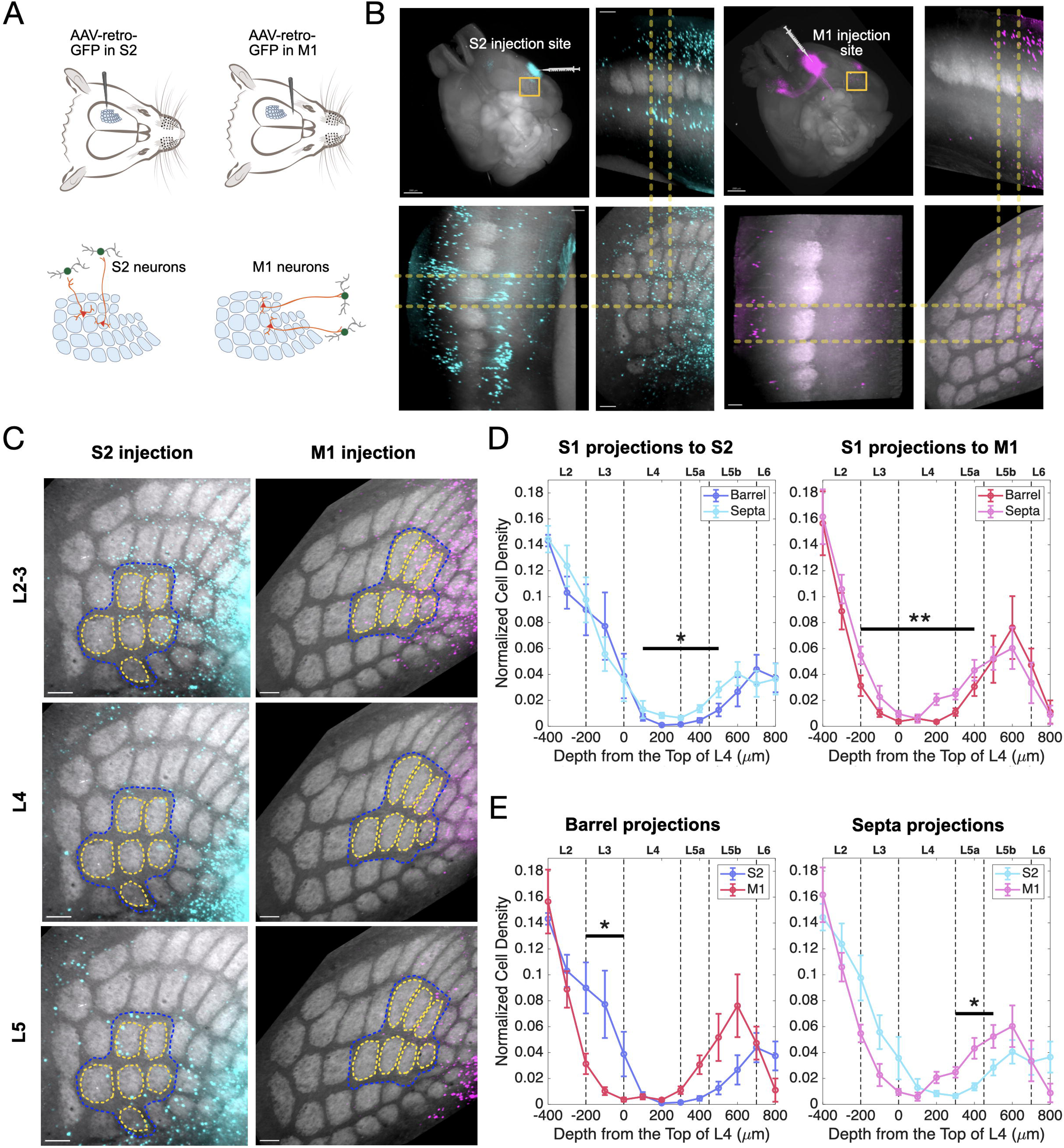
wS1 barrel and septa columns differentially project to wS2 and M1 (A) The schematics of the experimental protocol. The retro-AAV-CAG-GFP was injected in either S2, or M1 at P20-30 and the brains were dissected 4 weeks after the virus injection following the tissue clearing and whole brain imaging. (B) 3D reconstruction of the cleared brain with S2(Left panels) or M1(right panels) injection. X-Y, X-Z and Y-Z views are presented focus with a single barrel max projection. The brain was rotated to achieve the orientation that the septa is perpendicular to the X-Y view (top-down). Scale bars in whole brain view are set for 2000um and the scale bars in X-Y, X-Z and Y-Z views are set for 200um. (C) Max projection examples of S2-projected wS1 neurons and M1-projected wS1 neurons in L2-3, L4 and L5 in barrel vs septa. Quantified barrels (yellow lines) and septa (blue lines) are circled out which are perpendicular to X-Y (top-down) view. Scale bars are set for 200um. (D) Left: Normalized cell density profiles of S2 projection neurons at barrel and septa columns. 0µm represents the top of layer 4. L4 and L5a septa columns send significantly more projections to S2 than barrel column (N=4 mice) (p=0.0139, F=11.79, repeated-measures ANOVA). Right: Normalized cell density profiles of M1 projections neurons at barrel and septa columns. 0µm represents the top of layer 4. L3 and L4 septa columns send significantly more projections to M1 (N=6 mice) (p=0.0032, F=14.82, repeated measures ANOVA). (E) Left: Normalized cell density profiles of S2 and M1 projection neurons at barrel columns. 0µm represents the top of L4. L3 barrel columns send significantly more projections to S2 (N=4 mice with S2 injections, N=6 mice with M1 injections) (p=0.0293, F=7.02, repeated-measures ANOVA). Right: Normalized cell density profiles of S2 and M1 projections neurons at septa columns. 0µm represents the top of L4. L5a septal columns send significantly more projections to M1 N=4 mice with S2 injections, N=6 mice with M1 injections) (p=0.0402, F=5.98, repeated measures ANOVA).

As an internal control and in line with expectations, fewer projections arose from layer 4 barrels to either wS2 or M1 than from other cortical layers (Figure 6C and D). However, more neurons in septa between mid-L4 and upper-L5B sent wS2 projections (Figure 6D, left, turquois), and more neurons in septa from L3 to L4 project to M1, compared to the neurons in barrels (Figure 6D, right, magenta). Additionally, a secondary comparative analysis showed that barrels in L3 host significantly more wS2-projecting-neurons (Figure 6E, left, purple), while L5A septal domains host significantly more M1-projecting neurons (Figure 6E, right, magenta). This differential targeting suggests that the temporal whisker-evoked processing specialization of barrel and septa domains may shape how distinct streams of sensory information are conveyed to higher-order cortical areas, potentially influencing perceptual processing in wS2 and sensorimotor integration in M1.

## Discussion

In the absence of visual stimuli, humans explore objects through tactile interactions, where the initial touch begins the process of information accumulation, with subsequent touches refining the identification and characterization of the object. Rodents rely on a similar strategy during active whisking, where sequential whisker contacts allow for the progressive refinement of object recognition and spatial mapping (Armstrong-James and Fox, 1987; Kheradpezhouh et al., 2017; Petersen, 2007). This process involves parallel ascending pathways that transmit peripheral touch sensations to the cortex, with the primary somatosensory cortex mapping whisker spatial patterns directly onto cortical barrels, and in between spaces, septa (Van Der Loos and Woolsey, 1973; Woolsey et al., 1975).

Our study reveals that the functional segregation between barrel and septa domains in the mouse whisker primary somatosensory cortex (wS1) is not only anatomically defined by distinct thalamocortical projections but is also dynamically shaped by local inhibitory circuits, particularly those involving somatostatin-positive (SST+) interneurons.

Through a combination of functional and anatomical techniques, we show that the temporal divergence in spiking activity between barrel and septa domains is mediated by the short-term synaptic facilitation properties of SST+ interneurons and is critically dependent on the synaptic protein Elfn1. Because the narrow width of septa and the spatial sensitivity of multi-unit recordings preclude absolute cellular specificity, we interpret our electrophysiological measurements as reflecting septal-enriched versus barrel-enriched populations. Importantly, reanalysis using increasingly stringent spike-detection thresholds revealed a layer-specific dissociation: key domain-specific effects persisted selectively in Layer 4, whereas effects in L2/3 were attenuated. This threshold-dependent robustness supports the interpretation that the critical effects reported here arise from neurons spatially closer to the recording sites and are less influenced by probe orientation or distant sources, consistent with a local circuit origin in Layer 4.

Furthermore, using a decoder we show that this temporal divergence may affect the integration of sensory information during repeated whisker stimulation, which may be subsequently routed to differential downstream cortical areas, such as the secondary somatosensory cortex (wS2) and primary motor cortex (M1).

Among the inhibitory neuron classes, SST+ interneurons are known for their high spontaneous activity and powerful regulation of local neural networks through dense feedback connections onto nearby pyramidal neurons (Urban-Ciecko and Barth, 2016). In contrast, VIP+ interneurons influence network function through a disinhibitory circuit, inhibiting SST+ (and to a lesser extent PV+) interneurons upon activation, which in turn reduces inhibition on pyramidal cells (Karnani et al., 2016; Lee et al., 2013; Pi et al., 2013). Thus SST+ interneurons may contribute to barrel-septal identity using surround inhibition, while being regulated themselves by VIP+ interneuron activity under certain behaviorally relevant sensory-motor tasks. Additionally, fast-spiking (FS) interneurons, predominantly PV+ cells, form a distinct network electrically coupled with other FS cells and are preferentially activated by thalamocortical inputs, providing rapid feedforward inhibition that complements the slower, facilitating inhibition of SST+ networks (Gibson et al., 1999). Our analysis of SST+ cell distribution, shows a higher density in the septa in L4, compared to barrels. Interestingly, layer 4 SST+ interneuron axons remain within layer 4 (Xu et al., 2013) in the somatosensory cortex, unlike SST+ of layers 2/3 and 5/6 which are Martinotti type and target layer 1. In addition, while Martinotti cells receive facilitating synapses from pyramidal cells and form inhibitory synapses onto dendrites of neighboring pyramidal cells (Silberberg and Markram, 2007), non-Martinotti SST cells strongly inhibit fast spiking cells, but also stellate cells in layer 4 (Ma et al., 2006; Xu et al., 2013). During naturalistic whisking, SST+ and PV+ cells show divergent activation patterns, with FS cells showing transient excitation due to rapid synaptic depression, while SST+ cells show the opposite (Tan et al., 2008). These divergent dynamics are consistent with the presence of two parallel inhibitory networks in layer 4, where FS (PV+) cells exhibit strong short-term depression and LTS (SST+) cells display facilitation, enabling differential recruitment by temporal patterns of activity (Beierlein et al., 2003). It is currently unclear though under which behavioral circumstances SST+ cells of L4 would engage in inhibition of excitatory cells versus inhibition of PV+ cells. Our data suggest that under our experimental conditions, a bottom-up activation paradigm, SST+ cells overall engage in the progressive inhibition of the local L4 circuit, rather than disinhibition. Although we interpret divergence in L2/3 responses as potentially inherited from L4 dynamics, direct laminar recordings and manipulation would be required to confirm this. We propose that the divergence in MWS/SWS ratios across barrel and septal domains reflects dynamic microcircuit interactions, not fully or solely captured in SST+ density. Barrel domains, dominated by VPM inputs, engage PV+ and SST+ neurons to stabilize responses, with Elfn1-dependent facilitation gradually increasing inhibition during repetitive SWS. Septal domains, receiving facilitating POm inputs and having a higher L4 SST+ density, show progressive inhibitory buildup that amplifies the MWS/SWS ratio. Trans-laminar and lateral SST+ projections likely propagate these dynamics to L2/3, explaining divergence despite uniform SST+ density. Direct laminar cell-type specific and VPM/POm-specific optogenetic manipulations will be critical to test this model.

To assess whether temporal divergence in spiking carries meaningful sensory information, we conducted a detailed accumulative temporal decoding analysis. We find that in wild-type mice, decoding accuracy for SWS increased progressively with each additional pulse, particularly in the initial response window (1-50ms), indicating that domain-specific information accumulates primarily in early temporal segments during repeated whisker sampling. This progressive accumulation of decodable information was severely disrupted in Elfn1 KO mice, with cross-validation showing significantly reduced performance compared to wild-type, especially in early response segments. Interestingly, Elfn1 KO mice showed higher cross-validation performance in the later response segment (51-95ms), suggesting a temporal delay in information processing due to reduced progressive inhibition. Under multi-whisker stimulation, wild-type and knockout mice showed markedly different initial decoding accuracies, with the difference between accumulated and initial accuracies being lower in wild-type than in Elfn1 KO mice. Even though we have made significant efforts to define the columns in the barrel cortex, we are aware that the CLARITY-based passive clearing protocol leads to tissue expansion. In addition, different barrel columns were analyzed across different brains due to the technical difficulties, adding a small variability to the dataset. Regardless, in toto our anatomical connectivity mapping would suggest that disruption in temporal information processing in the two domains is crucial for maintaining the functional segregation of parallel processing streams to wS2 and M1.

The differential projection patterns of barrel and septa domains to wS2 and M1 further suggest that the temporal processing capabilities of these domains influence how sensory information is integrated with motor output. For example, the projection of septa domains to M1 may facilitate the rapid translation of sensory information into motor commands, while the projection of barrel domains to wS2 may support more detailed sensory discrimination. Our retrograde labeling data supports and expands on previous work proposing similar models (Alloway, 2008; Chakrabarti and Alloway, 2006), Further, this pathway-specific modulation is supported by evidence of learning-related plasticity in S1 cortico-cortical neurons, where M1-projecting neurons encode kinematic features and are involved in touch-related activity, while S2-projecting neurons enhance discrimination of trial types and decision-related patterns (Chen et al., 2015, 2013). Additionally, the preferential activation of inhibitory interneurons, such as SST+ neurons, by thalamocortical inputs, due to stronger excitatory synapses, underpins the initial processing stages in the barrel cortex (Cruikshank et al., 2007). The disruption of the informational integrity of these projections in Elfn1 KO mice could therefore have significant implications for sensorimotor integration and behavior, potentially impairing the adaptive plasticity observed in wild-type animals during task learning.

It is important to note that while our study focuses on the mouse barrel cortex, there are significant differences between rodent species, such as rats and mice, in terms of cortical organization and whisker system function. Most research on whisker stimulation had traditionally focused on rats (Estebanez et al., 2016; Petersen, 2007; Roy et al., 2011; Tanke et al., 2018). The existence of septa in the mouse barrel cortex has been debated, with some suggesting that the distinction between barrel and septal circuits is less pronounced in mice compared to rats (Bureau et al., 2006; Sato et al., 2007). For example, the mouse barrel cortex has fewer neurons per column and thinner septa compared to rats, which may influence the degree of functional segregation between barrel and septa domains. While many animals have whiskers, only a subset of animals have visible cortical barrels. For instance, cats have whiskers and brainstem barrelettes but lack cortical barrels (Nomura et al., 1986; Rice, 1985; Rice et al., 1985). Interestingly, the presence of barrels in the cortex seems to correlate with volitional whisking behavior, underlining the importance of one-to-one mapping for processing individual whisker information. Our findings show that septa are crucial in separating these parallel whisker streams by modulating the lateral cortico-cortical information flow during repetitive whisker touches. Our data support this modulation being significantly influenced by the short-term facilitation properties of synaptic inputs onto SST+ interneurons, facilitated by the Elfn1 gene, alongside distinct bottom-up inputs to each domain. The loss of temporal segregation in Elfn1 KO mice could hinder object recognition, which relies on multi-whisker integration in septa, while sparing simpler detection tasks mediated by barrels. Although Elfn1 is constitutively knocked out, this and earlier studies have found that barrel structure is preserved (Stachniak et al., 2023, 2019). Further, the distribution of Elfn1 expressing interneurons is not different in KO mice, suggesting minimal developmental disruption (Dolan and Mitchell, 2013). Nonetheless, we acknowledge that subtle circuit changes cannot be ruled out without the usage of time-depended conditional knockout of the gene.

In summary, our study demonstrates that there exists a functional segregation of barrel and septa domains in the mouse whisker somatosensory cortex, and that it is dynamically shaped by the temporal dynamics of SST+ interneuron activity, imparted by the synaptic protein Elfn1. This temporal divergence is essential for the integration of sensory information over repeated whisker deflections and for the differential projection-dependent information processing in downstream cortical areas. These findings highlight the importance of local inhibitory circuits in shaping cortical processing and offer new insights into mechanisms underlying sensory perception and behavior.

## Supporting information

Response to Reviewers

## Acknowledgments

We thank members of the Karayannis lab for suggestions and comments on the manuscript. Authors also thank George Kanatouris for helping with animal handling and breeding.

## Funding

This work was supported by Swiss National Science Foundation Grant S-41260-01-01 (T.K) and European Research Council Grant 679175 (T.K).

## Author Contributions

A.Ö.A. curated the data, analyzed in-vivo silicon probe recordings and the two-photon calcium imaging data, performed the decoder analysis and wrote the manuscript. T.J.S. conceptualized the study, maintained Elfn1 KO line, performed earlier electrophysiological characterizations and contributed to the drafting of the manuscript, J.W.Y. performed silicon probe recordings, AvdB collected two-photon calcium imaging data, L.C. performed AAV virus injection, tissue clearing and light-sheet data acquisition and analysis, R.K. performed tissue clearing and light-sheet image acquisition, T.K. conceptualized and supervised the study and wrote the manuscript.

## Declaration of Interest

The authors declare no competing interests.

## Methods

### RESOURCE AVAILABILITY

#### Lead Contact

Further information and requests for resources and reagents should be directed to the Lead Contact, T.K.

#### Materials Availability

This study did not generate new unique reagents.

#### Data and Code Availability

The datasets and analysis routines are available from the first author on reasonable request.

### Mice

All the animal experiments followed the guidelines of the Veterinary Office of Switzerland and were approved by the Cantonal Veterinary Office Zurich and the University of Zurich. husbandry with a 12-h reverse dark-light cycle (7 a.m. to 7 p.m. dark) at 24 °C and variable humidity. Adult (5- to 10-week-old) male C57BL6J wide type mice were used for retrograde tracing. Animal lines used in this study are VIP-IRES-Cre (Vip^tm1(cre)Zjh^/J)(Taniguchi et al., 2011), SST-IRES-Cre (Sst^tm2.1(cre)Zjh^/J) (Taniguchi et al., 2011), Ai14 (B6;129S6-Gt(ROSA)26Sor^tm14(CAG-tdTomato)Hze^/J)(Madisen et al., 2010) and Elfn1 KO (Elfn1tm1(KOMP)Vlcg). Both male and female mice were used in all experiments.

### In-Vivo Silicon Probe Recording

We used 4 Elfn1 KO and 4 WT littermates at the age of P20 to P30 for the multi-electrode silicon probe recordings. Mice were under urethane induced anesthesia (1.5g/1kg) throughout the experiment. A heating pad was used to maintain the mouse’s body temperature at 37°C. The depth of anesthesia was checked with breathing speed and paw reflexes throughout the experiment. The skull of the right hemisphere was exposed by removing the skin on top, and a metallic head holder was implanted on the skull with cyanoacrylate glue and dental cement. A 26G needle was used to open a ∼2mm x 2mm cranial window which exposed the S1 barrel field (wS1). Extreme care was taken not to cause damage or surface bleeding during surgery.

wS1 neural activities were recorded with an 8-shank-64-channel silicon probe. Each of the 8 shanks has 8 recording sites (100μm apart). The distance between each shank is 200μm (A8x8-Edge-5mm-100-200-177-A64, NeuroNexus Technologies, Ann Arbor, MI, USA). The silicon probe was labeled with DiI (1,1′-dioctadecyl-3,3,3′,3′-tetramethylindocarbocyanine, Molecular Probes, Eugene, OR, USA) and inserted perpendicularly into the barrel cortex. A silver wire was placed into the cerebellum as a ground electrode. All data were acquired at 20 kHz and stored with MC Rack (RRID:SCR_014955) software (Multi Channel systems). The total duration of multi-electrode recordings varied between 3 to 5 hours. After each experiment, the animal was deeply anesthetized by ketamine (120 mg/kg, ketamine, 50 mg/mL, HamelnPharma, Hameln Germany) and perfused through the aorta with ringer solution. The brain was kept in 4% PFA. Tangential sections (200-μm thick) were prepared for vesicular glutamate transporter 2 (vGlut2, Synaptic Systems, 135404; 1:1000) immunohistochemistry. By combining the DiI and vGlut2 staining, the insertion position of the 8 shank probes was identified in the barrel cortex. DiI marks contained within the vGlut2 staining were defined as barrel recordings, while DiI marks outside vGlut2 staining were septal recordings.

### Whisker Stimulation

Either a single principal whisker (SW, single whisker) or multiple whiskers (MW, multi whisker) were stimulated 1mm from the snout in rostral-to caudal direction (about 1mm displacement) using a stainless-steel rod (1 mm diameter) connected to a miniature solenoid actuator. The movement of the tip of the stimulator bar was measured precisely using a laser micrometer (MX series, Metralight, CA, USA) with a 2500 Hz sampling rate. The stimulus takes 26ms to reach the maximal 1 mm whisker displacement, with a total duration of 60ms until it reaches baseline (Yang et al., 2017). Both SW and MW stimulation was performed at 10Hz for 2s and each stimulation repeated for 20 times (20 Trials). The inter-trial interval was 30s. In every experiment, multiple whiskers were singly stimulated to increase the probability of finding a proximal barrel-septa and neighboring barrel triplets. We first performed multi-whisker stimulation and at the end individual single whisker stimulations. Individual whisker stimulation was performed by attaching a pipet tip to the stimulator, which was then placed in contact with the principal whisker under a dissecting stereoscope. We trimmed the whiskers where necessary, to avoid them touching each other and to avoid stimulating multiple whiskers. By putting the pipette tip very close (almost touching) to the principal whisker, the movement of the tip (limited to 1mm) reliably moved the principal whisker, as observed under the stereoscope. MW stimulation was performed by contacting multiple whiskers with 1cm-wide masking tape attached to the pipet tip.

### In vivo two-photon calcium imaging

Neuronal ensembles in superficial layers of the principal whisker barrel field mapped by intrinsic signal imaging were bolus-loaded with the AM ester form of Oregon Green BAPTA-1 by pressure injection (OGB-1; 1 mM solution in Ca^2+^-free Ringer’s solution; 2-min injection at 150–200 µm depth) as described previously (Stosiek et al., 2003). The craniotomy was then filled with agarose (type III-A, 1% in Ringer’s solution; Sigma) and covered with an immobilized glass plate. Two-photon Ca^2+^ imaging was performed with a Scientifica HyperScope two-photon laser scanning microscope one hour after bolus loading using a Ti:sapphire laser system at 900 nm excitation (Coherent Chameleon; ∼120 femtosecond laser pulses). Two-channel fluorescence images of 256 × 128 pixels at 11.25 Hz (HyperScope galvo-mode) were collected with a 16x water-immersion objective lens (Nikon, NA 0.8). In all, 3–5 separate spots have been imaged per animal. Per imaging spot, 10 trials of 20s-long evoked activity recorded for each of single- and multi- whisker stimulation paradigms. Data acquisition was controlled by ScanImage (RRID:SCR_014307) (Pologruto et al., 2003).

### Analysis of calcium imaging data

Ca^2+^ imaging data is analyzed using custom MATLAB scripts. First, fluorescence image time-series for a given region were concatenated. The concatenated data was then aligned using a Fourier domain cross-correlation based subpixel registration algorithm (Guizar-Sicairos et al., 2008) to correct for translational drift (registered on red tdTomato channel and transferred to OGB-1 channel). Average intensity projections of the imaging data were used as reference images to manually annotate regions of interest (ROIs) corresponding to individual neurons. Neurons with somata partly out-of-focus were not included. Ca^2+^ signals were expressed as the mean pixel value of the relative fluorescence change Δ*F/F* = (*F*-*F_0_*)/*F_0_* in each given ROI. *F*_0_ was calculated as the bottom 5% of the fluorescence trace. Neuropil patches surrounding each neuron is defined by all pixels not assigned to a neuronal soma or astrocyte of the corresponding neuron ROI annotation (Peron et al., 2015) (A disk shaped region around the neuron of interest with excluding any intersecting neighboring neuronal ROI). Neuropil correction is performed as *F*_corrected_ = *F*_neuron_ – *α*F*_neuropil_. α is estimated for each imaging spot separately using the formula *F*_blood_vessel_/*F*_surrounding_neuropil_ (Kerlin et al., 2010). For each stimulus, the evoked responses of 10 trials were analyzed and the response magnitude expressed as the mean of the evoked Δ*F/F* integral (%·s; integral of the first 2s response starting at stimulus onset).

### Analysis of Silicon Probe Data

Extracellular silicon probe data were analyzed using a custom-made Matlab script (MATLAB (RRID:SCR_001622), Version: 2024a, MathWorks, MA, USA). The raw data signal was band-pass filtered (0.8-5 kHz), and the multi-unit activity (MUA) was extracted with the threshold of 7.5 times the standard deviation (SD) of baseline (Yang et al., 2017). The current source density (CSD) map was used to identify laminar probe contact location. The earliest CSD sink was identified as layer 4, followed by L2-3 (Reyes-Puerta et al., 2015; Van Der Bourg et al., 2017). Post hoc histology was then performed on plane-aligned brain sections which would allow us to detect barrels and septa, so as to confirm the insertion domains of each recorded shank. Layer specificity of each electrode could therefore not be confirmed by histology as we did not have coronal sections in which to measure electrode depth. Instead, the recording sites above the one in L4 were designated as 2/3 and the ones below as deeper layer ones. The analysis of the MUA activity was smoothed using a Gaussian kernel (0 mean, 5ms sigma) and used for also delineating the different layers.

The Area Under the Curve (AUC) was computed as the integral of the smoothed firing rate (spikes per millisecond) over a 50ms window following each whisker stimulation pulse, using trapezoidal integration. Firing rate data for layer 4 barrel and septal regions in wild-type (WT) and knockout (KO) mice were smoothed with a 3-point moving average and averaged across blocks of 20 trials. Plotted values represent the percentage ratio of multi-whisker (MW) to single whisker (SW) AUC with error bars showing the standard error of the mean. Each data point reflects the mean AUC ratio for a stimulation pulse across approximately 11 blocks (220 trials total). The y-axis indicates percentages.

### Decoder Analysis

Recordings from the principal barrel, adjacent septa, and a neighboring unstimulated barrel were organized into three matrices for wild-type (WT) animals: a 280 × 95 × 20 matrix for the stimulated barrel (14 Barrels, 95ms, 20 pulses), a 180 × 95 × 20 matrix for the septa (9 Septa, 95ms, 20 pulses), and a 360 × 95 × 20 matrix for the neighboring barrel (18 Neighboring barrels, 95ms, 20 pulses). For Elfn1-KO animals: 11-barrel columns, 7 septal columns, 11 unstimulated neighbors from N=4 mice.

Each matrix’s dimensions correspond to the number of recording channels, time points (95ms epochs), and 20 individual stimulation trials, respectively. Although each trial originally spanned 2000ms (20 pulses at 10Hz), the analysis focused on the first 95ms post-stimulation (95ms instead of 100ms). To examine the impact of different temporal windows on classification performance, three versions of the analysis were performed by varying the time-range parameter, 1 to 95ms for the full epoch, 1 to 50ms for the first part, and 51 to 95ms for the second part.

For each stimulation trial, data was accumulated pulse-by-pulse. In each iteration (ranging from 1 to 20), the corresponding segments from the three matrices were reshaped into two-dimensional feature matrices by concatenating the selected time points across the accumulated pulses. Labels were then assigned to the reshaped data (‘B’ for the barrel, ‘S’ for septa, and ‘N’ for the neighboring barrel).

We employed a classification approach utilizing error-correcting output codes (ECOC) models with a gentle adaptive boosting (GentleBoost) strategy to train classifiers. This method is particularly effective in handling imbalanced data and unequal misclassification costs. Like Logit Boost, each weak learner in the ensemble fitted a regression model to response values, 𝑦_𝑛_ ∈ {–1, +1}. The mean-squared error is computed as:

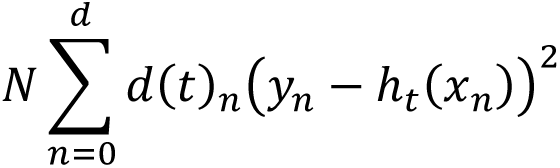

where 𝑑(𝑡)_𝑛_ represents observation weights at step t (summing up to 1) and ℎ_𝑡_(𝑥_𝑛_) denotes predictions from the regression model ℎ_𝑡_ fitted to response values 𝑦_𝑛_. As the strength of individual learners diminishes, the weighted mean-squared error converges towards 1.

Three distinct evaluations were performed. WT-to-KO Prediction, the ECOC classifier trained on WT data (compiled from the three regions) was used to predict labels from corresponding recordings in knockout (KO) mice. Within-Group Cross-Validation (WT), A cross-validation procedure was applied to the WT dataset using MATLAB’s *crossval* function to assess classifier performance internally. Within-Group Cross-Validation (KO), Similarly, an ECOC classifier was trained and cross-validated on the KO dataset.

For each evaluation, confusion matrices were generated and summary accuracy statistics computed. Accuracy was tracked as a function of the number of accumulated pulses (from 1 to 20). The accuracy value represents the overall classification accuracy, defined as the proportion of correctly predicted samples across all classes. Specifically, it is calculated as the sum of true positives (TP) for all classes obtained from the trace of the confusion matrix (i.e., the diagonal elements where predicted and actual labels match) divided by the total number of samples. The formula used is:

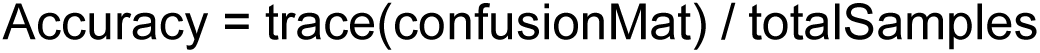

where totalSamples is the sum of all entries in the confusion matrix.

This metric reflects the overall correctness of the classifier and is naturally constrained between 0 and 1. The function first constructs the confusion matrix confusionMat, where rows represent actual classes and columns represent predicted classes, either by computing it from labels using MATLAB’s confusionmat or using a precomputed matrix. The number of classes is determined from the matrix dimensions, and the total number of samples is the sum of all matrix entries. For each class, the function calculates TP (correct predictions for that class), TN (correct predictions for all other classes), FP (false positives for that class), and FN (false negatives for that class) by manipulating the confusion matrix.

To characterize the evolution of classification accuracy with increasing pulse number, non-linear exponential functions of the form:

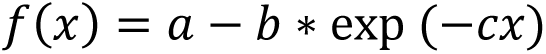

were fitted to the accuracy data using a non-linear least squares approach (implemented via the *fminspleas* function). Separate fits were obtained for the WT-to-KO prediction, WT cross-validation, and KO cross-validation accuracy curves. The fitted curves and raw accuracy data were then plotted. This integrated analysis pipeline allowed for systematic comparison of classification performance across spatial locations and conditions and an assessment of the temporal dynamics underlying the neural responses to single-whisker stimulation. The same strategy has been used for multi-whisker decoding but between only Barrel and Septa.

### Retrograde tracing

Mice aged from P10-37 were anesthetized by isoflurane and a small craniotomy was made above the injection site (M1 or S2). 200nl retrograde AAV virus, ssAAV-retro/2-CAG-EGFP-WPRE-SV40p(A) (physical titer: 5x10^12^ vg/ml) was injected 200-500µm deep in the M1 or S2. Virus was purchased from Zurich Viral Vector Facility (Cat#: V24-retro). During the retrograde virus injection, the location of M1 and S2 injections was determined by stereotaxic coordinates (Yamashita et al., 2018). After acquiring the light-sheet images, we were able to post hoc examine the injection site in 3D and confirm that the injections were successful in targeting the regions intended. Although it would have been informative to do so, we did not functionally determine the whisker-related M1 and whisker-related S2 region in this experiment.

### Tissue Processing for Passive Clearing and Imaging

The method used for hydrogel-based tissue clearing is explained in detail elsewhere (Chung and Deisseroth, 2013). Briefly, Mice age P29-70 (4 weeks after retrograde AAV virus injection) or SST-Ai14 and VIP-Ai14 mice age from P14-77 were transcardially perfused using 1x PBS and Hydrogel solution (1% PFA, 4% Acrylamide, 0.05% Bis). The collected brains were post-fixed for 48 h in a Hydrogel solution (1% PFA, 4% Acrylamide, 0.05% Bis). Afterward, the Hydrogel polymerization was induced at 37 °C. Following the polymerization, the brains were immersed in 40 mL of 8% SDS and kept shaking at room temperature until the tissue was cleared sufficiently (15-90 days depending on the animal’s age). Finally, after 2–4 washes in PBS, the brains were put into a self-made refractive index matching solution (RIMS). They were left to equilibrate in 5 mL of RIMS for at least 4 days at RT (Room Temperature) before being imaged. After clearing, brains were attached to a small weight and loaded into a quartz cuvette, then submerged in RIMS, and imaged using a home-built mesoscale selective plane illumination microscope (mesoSPIM, Voigt et al., 2019). The microscope consists of a dual-sided excitation path using a fiber-coupled multiline laser combiner (405, 488, 515, 561, 594, 647 nm, Omicron SOLE-6) and a detection path comprising an Olympus MVX-10 zoom macroscope with a 1× objective (Olympus MVPLAPO 1x), a filter wheel (Ludl 96A350), and a scientific CMOS (sCMOS) camera (Hamamatsu Orca Flash 4.0 V3). For imaging tdTomato and eGFP a 561 nm excitation with a 594 long-pass filter and 488 nm & 520/35 (BrightLine HC, AHF) were used, respectively. The excitation paths also contain galvo scanners (GCM-2280-1500, Citizen Chiba) for light-sheet generation and reduction of streaking artifacts due to absorption of the light-sheet. In addition, the beam waist is scanning using electrically tunable lenses (ETL, Optotune EL-16-40-5D-TC-L) synchronized with the rolling shutter of the sCMOS camera. This axially scanned light-sheet mode (ASLM) leads to a uniform axial resolution across the field-of-view of 5–10 µm (depending on zoom and wavelength). The magnification 0.8X (pixel size: 8.23 µm) was used for the view of the whole brain and 2X (pixel size: 3.26 µm) was used for the view of barrel cortex. Further technical details of the mesoSPIM are described elsewhere (Vladimirov et al., 2023; Voigt et al., 2019).

### Analysis of cleared brain imaging data

Barrel cortex was imaged in 3D with 2X objective. The images were processed through Fiji and Imaris to visualize the 3D structure of the barrel cortex. XY-, YZ-, XZ- projections were visualized in Imaris, which allows us to adjust the correct angle for each barrel column to be vertical against XY-plane. This method allowed us identify barrel and septa in layers 2-3 and 5 and layer 4. In addition, the depth information is also precisely acquired for each barrel column. SST+ or VIP+ neuron numbers, S2-projected and M1-projected S1 neuron numbers were counted in icy. Max projection for each 50 ums along barrel columns was acquired and neurons on each max projection image were counted automatically by icy (RRID:SCR_010587) with the same threshold for all the brains. Each barrel was drawn according to the barrel map from each brain by max projection. The area of septa was calculated by subtracting the whole selected area from all the selected barrel areas.

### t-SNE Analysis

To visualize the segregation of neural responses across the different cortical domains, we applied t-distributed Stochastic Neighbor Embedding (t-SNE) to the averaged datasets. t-SNE was used to reduce the high-dimensional data into a 2D space for visualization. The parameters for t-SNE were set to use the *barneshut* algorithm for efficiency, with a perplexity of 9 and exaggeration of 3 to enhance cluster separation. The resulting t-SNE coordinates were plotted using MATLAB’s *gscatter* function to distinguish between the different domains (Barrel, Septa, Neighboring). This analysis allowed us to visually inspect the clustering of neural responses from different cortical domains.

In summary, each point in the t-SNE plots represents an averaged response across 20 trials for a specific domain (barrel, septa, or neighbor) and genotype (WT or KO), with approximately 14 points per domain derived from the 280 trials in each dataset. The input features are preprocessed by averaging blocks of 20 trials into 1900-dimensional vectors (95ms × 20), which are then reduced to 2D using t-SNE with the specified parameters. This approach effectively highlights the segregation and clustering patterns of neural responses across cortical domains in both WT and KO conditions.

### Statistical Analysis of the Multi-Unit Activity

To evaluate the statistical significance of differences in multi-unit activity (MUA) responses between barrel (B), septa (S), and neighboring barrel (N) domains across 20 pulses of whisker stimulation, we performed a pulse-by-pulse comparison using the non-parametric Mann-Whitney U test (also known as the Wilcoxon rank-sum test). This analysis was conducted on the averaged MUA data recorded from Layer 4 of the whisker somatosensory cortex (wS1) in wild-type (WT) and Elfn1 knockout (KO) mice during single-whisker stimulation (SWS) and multi-whisker stimulation (MWS) paradigms, as detailed above. For each of the 20 stimulation pulses (1 to 20), the MUA data for each domain were reshaped and averaged across trials. The Mann-Whitney U test was then applied pairwise to assess differences between domains: barrel vs. septa (B vs S), barrel vs. neighboring barrel (B vs N), and septa vs. neighboring barrel (S vs N). The resulting p-values were stored in a 3×20 matrix, where rows correspond to the pairwise comparisons and columns represent the pulse number. To visualize the statistical outcomes, a heatmap was generated using MATLAB (Version 2024a, MathWorks, MA, USA). A custom discrete colormap was created by adapting the hot colormap, which originally provides a gradient of 10 colors ranging from black (low values) to white (high values). We selected specific indices from this gradient (10, 8, 6, 4, 2, 1) to define a 10-level discrete colormap, assigning the darkest shade (black) to p-values ≤ 0.05 (indicating statistical significance) and progressively lighter shades to higher p-values, with the lightest shade (white) representing p-values > 0.05. The colormap was inverted to ensure that significant p-values (≤ 0.05) appeared as light shades against a dark background, enhancing visual contrast. The heatmap was plotted with p-values scaled between 0 and 0.05, using the custom colormap.

### Statistical Analysis

Data are represented as mean ± SEM unless stated otherwise. Statistical comparisons have been done using a one-tailed Mann-Whitney-U test for non-paired data and one-tailed Wilcoxon signed-rank test for paired comparisons (the statistical significance threshold was set to p<0.05; in the Figures, different degrees of evidence against the null hypothesis are indicated by asterisks (p<0.05: *; p<0.01: **; p<0.001: ***). Cell densities in Figures 2 and 6 were compared using repeated measures ANOVA. All tests were conducted using custom codes in Matlab.

## Figure Legends

**Supplementary Figure 1.**
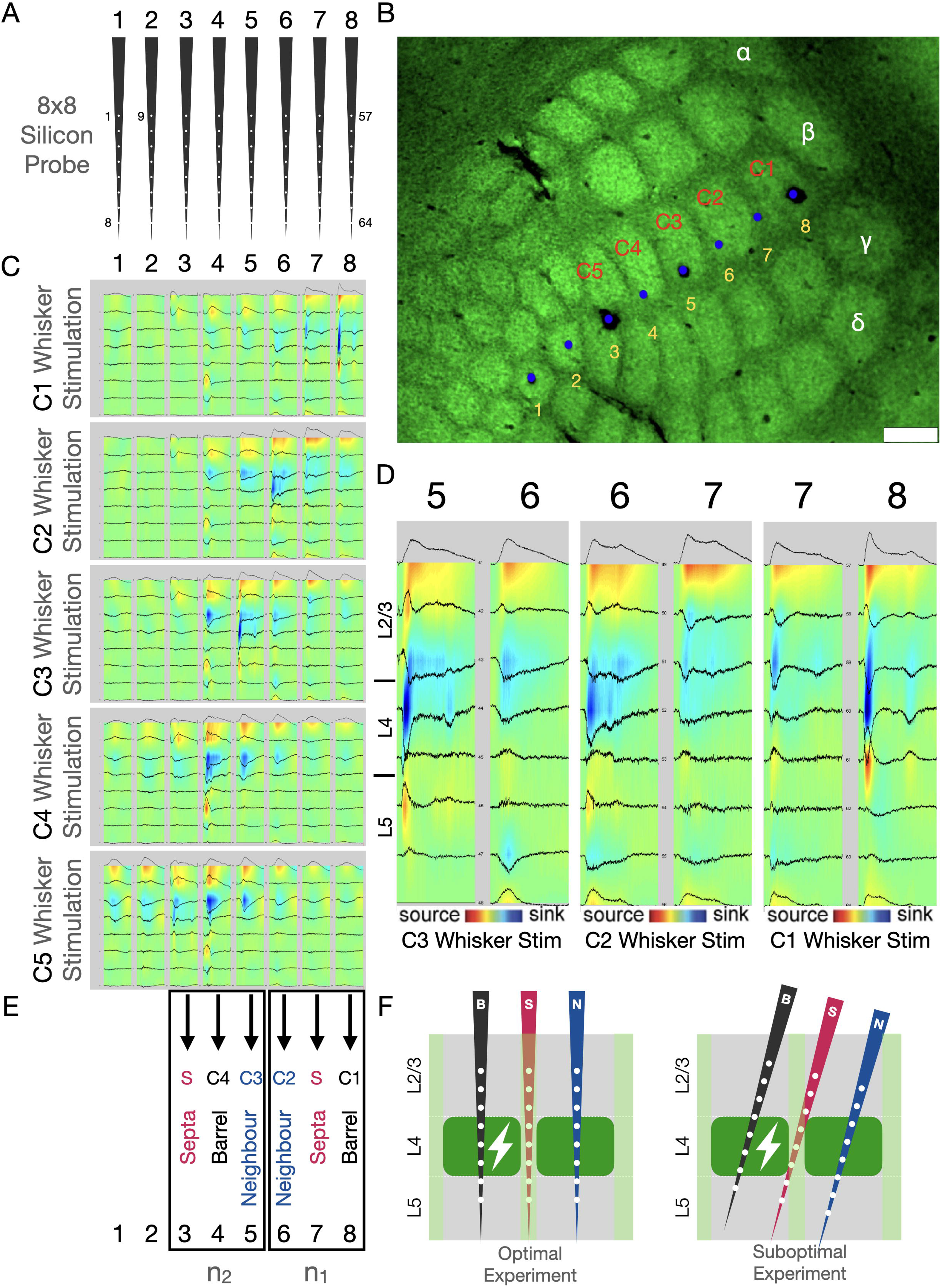
An example of assignment of barrel, septa and Neighbor electrodes using histology and current source density (CSD) analysis. (A) Shank electrode map of 8x8 silicon probes. (B) Probe locations were labeled (DiI) after each experiment using histology (vGlut2 staining). Matching shank locations are indicated using yellow numbers. Blue dots represent the insertion sites. The red and white text depicts corresponding barrel names. (Scale bar: 250µm) (C) All whiskers are stimulated one-by-one. C1 to C5 barrel responses upon single whisker stimulation are shown from top to bottom. (D) A closer look on C1, C2 and C3 whisker stimulations and corresponding CSD at shanks 5,6,7 and 8. (E), Barrel (B), adjacent septa (S) and the adjacent neighboring barrel (N) were assigned according to the probe insertion sites and L4 barrel autofluorescence in combination with CSD responses. For example, while shanks 6 and 8 show strong response upon single whisker stimulation (and hence assigned as barrels), shank 7 does not show the expected response (hence assigned as septa). Only the triplets of electrodes were used in the analysis in which principal barrel (B), adjacent septa (S) and the adjacent neighboring barrel (N) were captured by the probe insertion sites for any experiment (Left). 14-barrel columns, 9 septal columns, 18 unstimulated neighboring barrels from N=4 mice. (F) An ideal schematic representation (on the left). Realistically, we cannot always be sure if an entire shank is consistently going through a barrel or septal column due to the curvature of the cortex (on the right) for above and below of layer4.

**Supplementary Figure 2.**
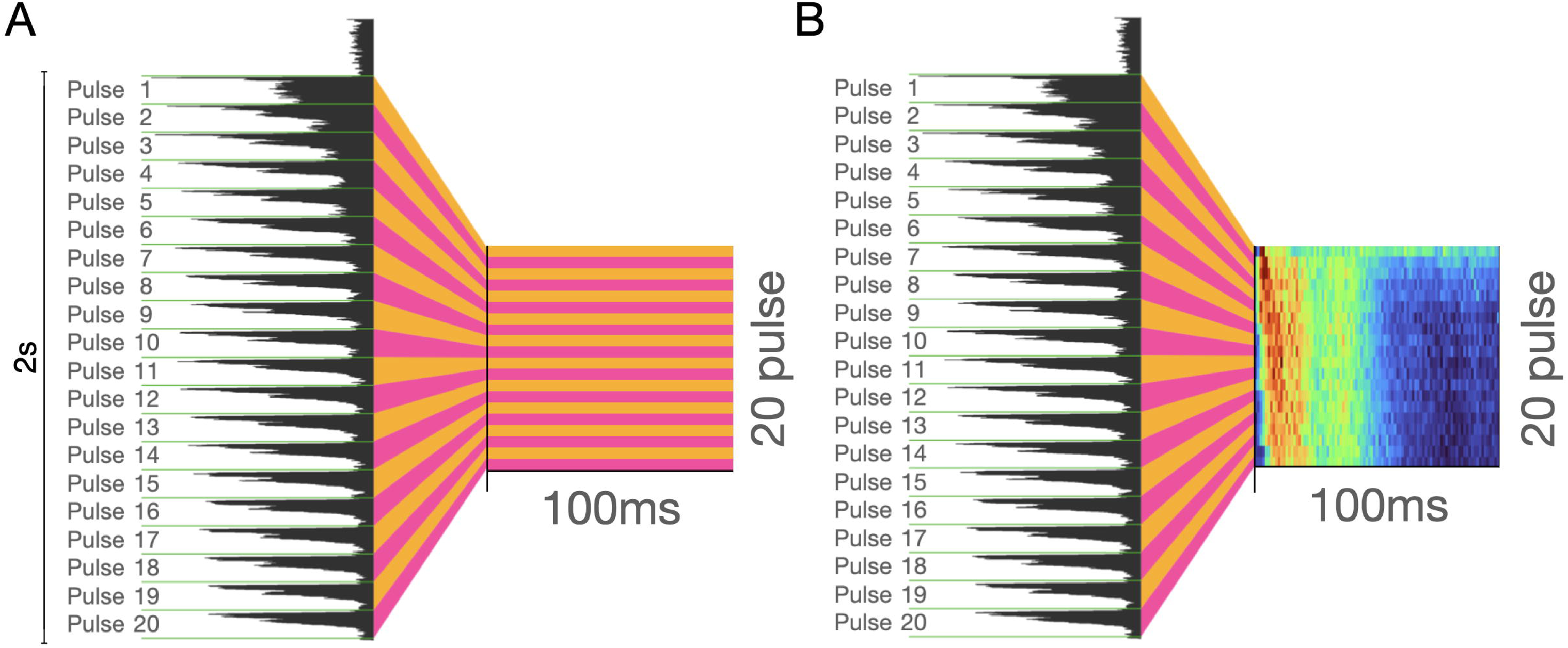
Representation of the conversion of the two-second-long single trial recordings into 20x100ms matrices. (A) Schematic representation. (B) An example data. The last 5ms of each pulse have been discarded in further analysis.

**Supplementary Figure 3.**
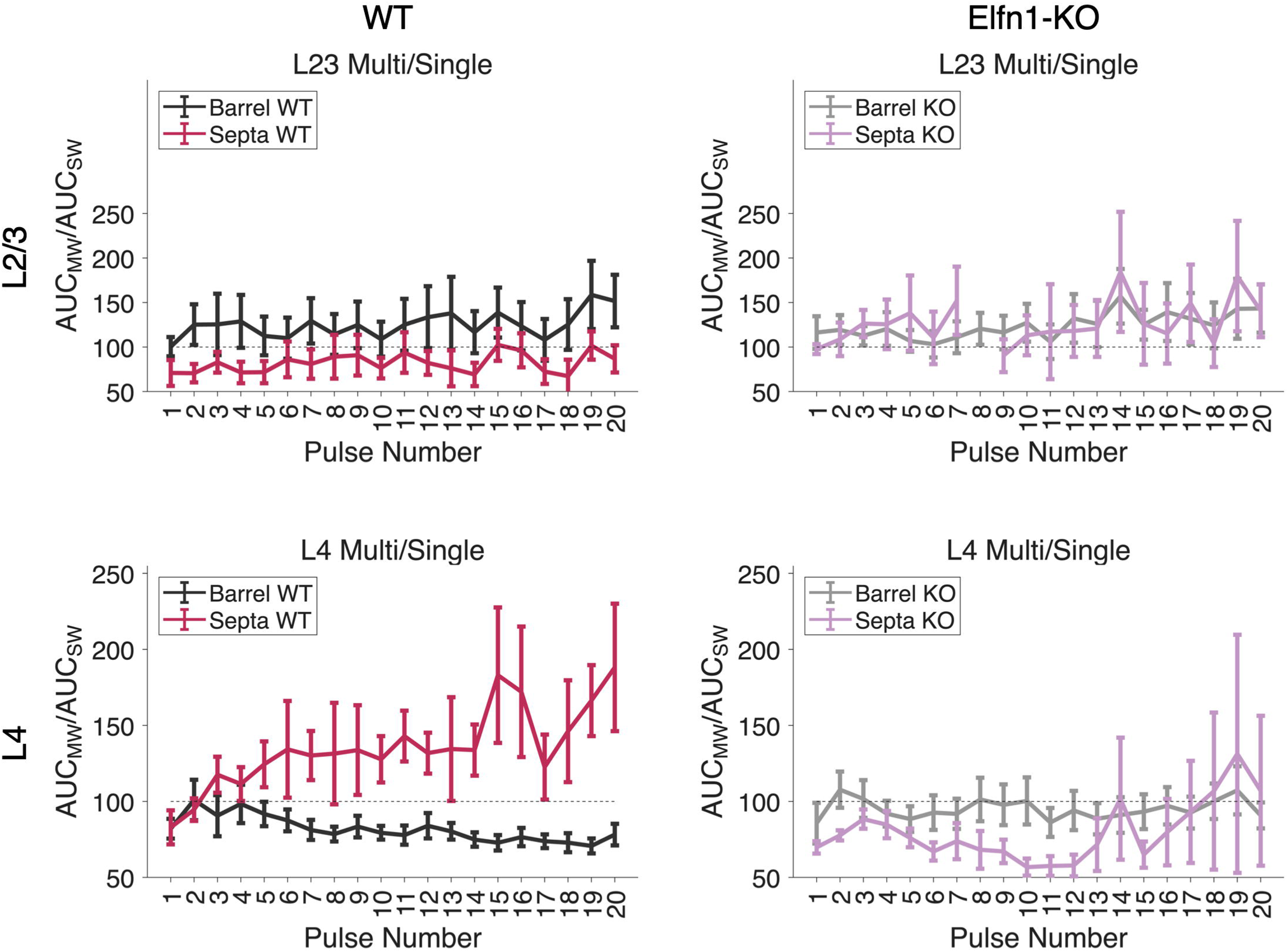
Re-analysis of Layer 2/3 (Top Row) and Layer 4 (Bottom Row) MW/SW data with a more stringent spike detection threshold of SD>9.5.

**Supplementary Figure 4.**
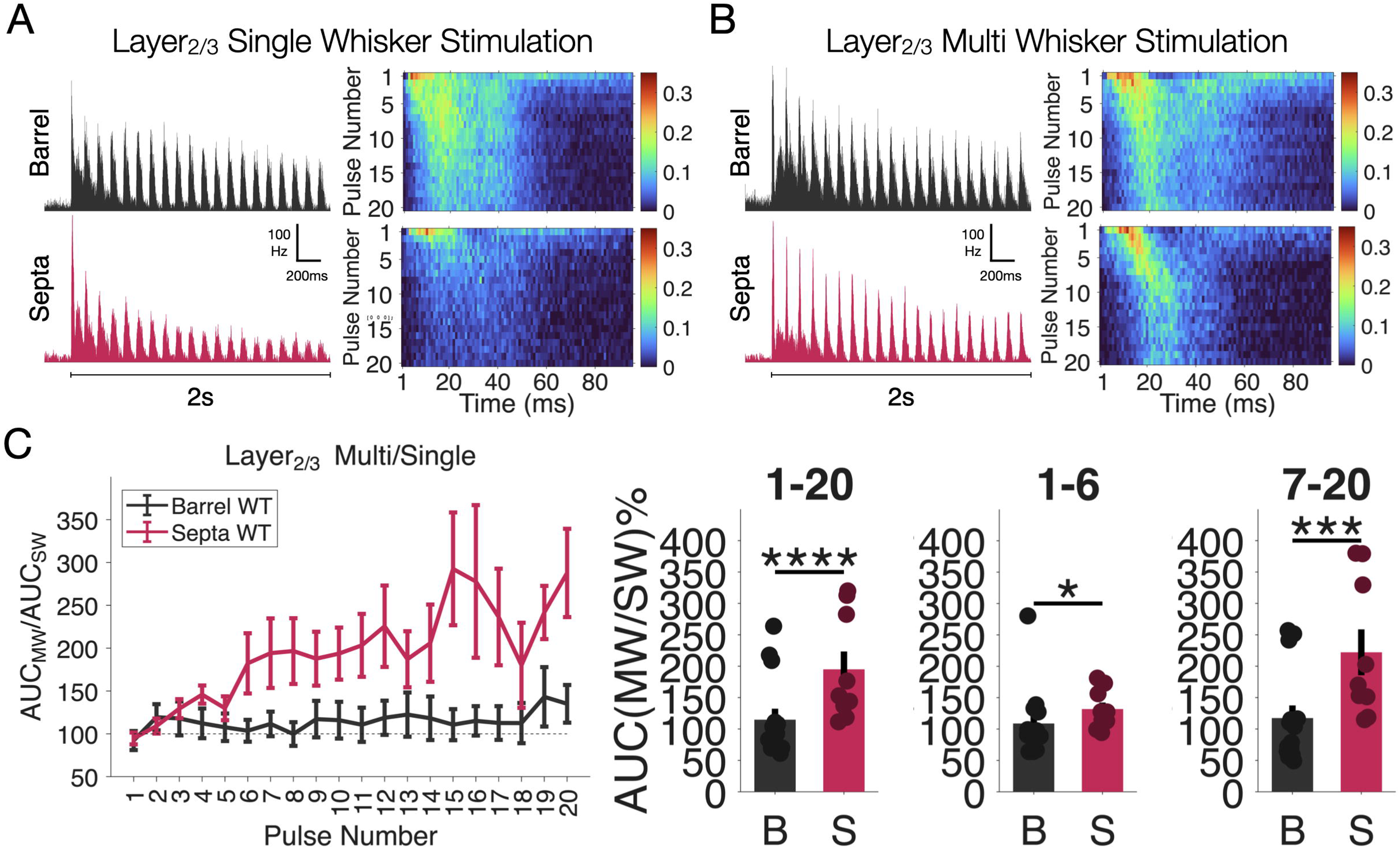
Divergence between barrel and septa domains upon repeated whisker stimulation is also found in Layer 2/3 of mouse wS1. (A) Left: Average multi-unit firing profiles recorded at Layer 2/3 of barrel (black), unstimulated neighbor (blue) and septa (pink) columns upon 2s-long 10Hz repeated single whisker stimulation. Right: Another representation of the same data but converted into an image by representing each stimulation pulse response on one row. (B) Left: Average multi-unit firing profiles recorded at Layer 2/3 of barrel (black), unstimulated neighbor (blue) and septa (pink) columns upon 2s-long 10Hz repeated multi whisker stimulation. Right: Another representation of the same data but converted into an image by representing each stimulation pulse response on one row. (C) Left: Temporal ratio dynamics (average AUC MW responses divided by average AUC SW responses) for barrel and septa for layer2/3. Right: statistical comparison of average AUC ratios of all pulses (1-20), first six (1-6) or last 14 pulses (7-20), respectively (stars represent *: p<0.05; **: p<0.01; ***: p<0.001 in F and G. Wilcoxon rank sum test).

**Supplementary Figure 5.**
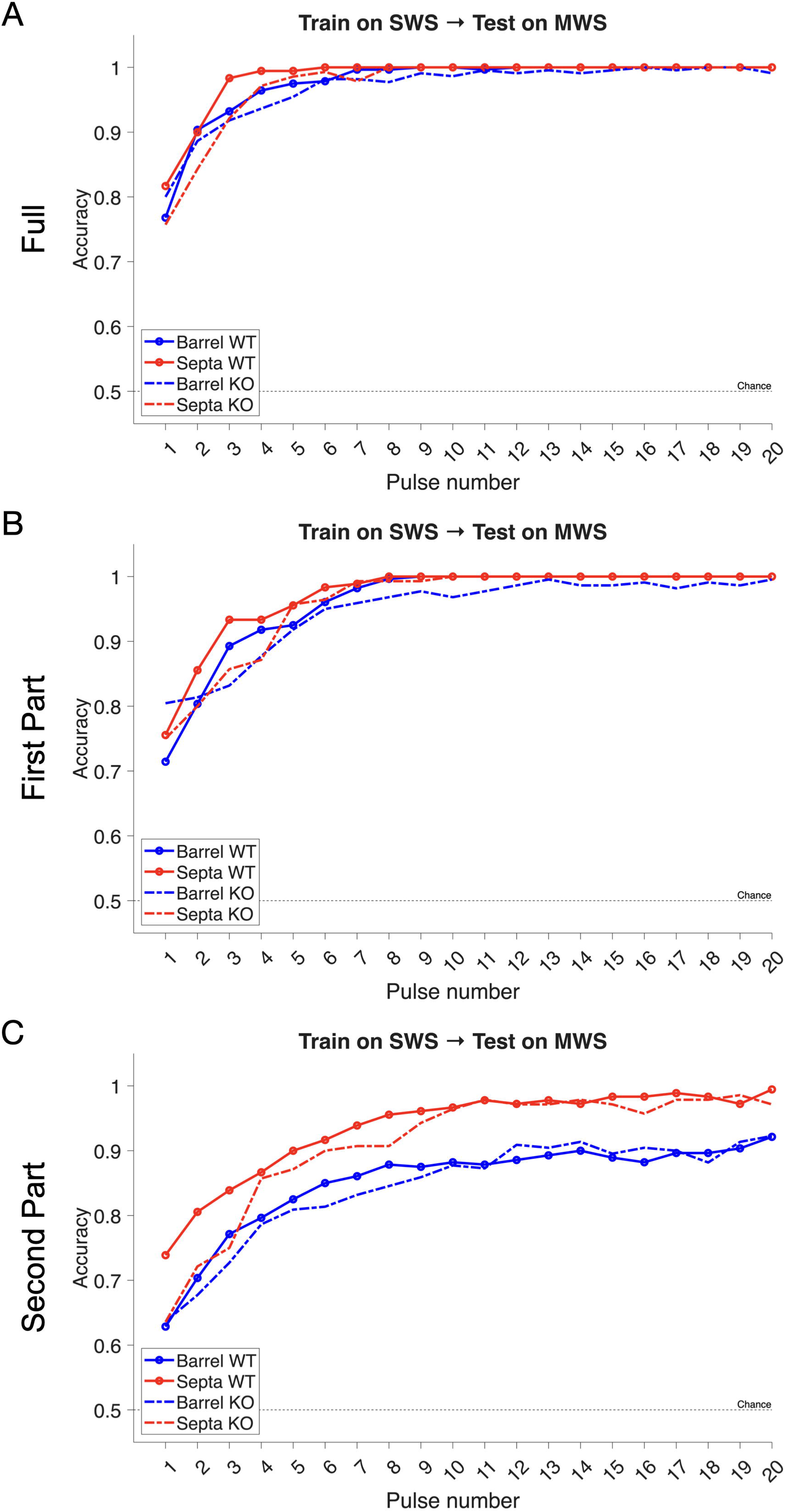
Cross-condition decoding between single-whisker (SWS) and multi-whisker stimulation (MWS). Recordings from the principal barrel and adjacent septa were organized into two matrices for two separate WT and KO decoders. To assess generalization, decoders were trained on MWS responses and tested on SWS responses. Data were accumulated pulse-by-pulse, and in each iteration (1–20) the corresponding trial segments were concatenated across pulses to construct feature matrices. Decoding performance was computed for three temporal windows: (A) the full response (1–95ms); (B) the first epoch (1–50ms), and (C) the second epoch (51–95ms).

## Notes

### Competing Interest Statement

The authors have declared no competing interest.

### Summary of Updates

This revised version of the manuscript incorporates conceptual clarifications, additional analyses, and textual revisions in response to reviewer feedback, with the goal of strengthening interpretation while preserving the scope of the experimental data. First, we clarified the interpretation of in vivo multi unit activity recordings obtained from barrel and septal domains. Given the narrow width of septa and the spatial sensitivity of multi unit recordings, we now explicitly describe septal recordings as reflecting septal enriched populations rather than exclusively septal neurons. To directly address concerns about spatial specificity, we performed additional analyses using a more stringent spike detection threshold. These analyses revealed a layer specific dissociation, with domain specific multi whisker versus single whisker response differences persisting selectively in Layer 4 while effects in Layer 2/3 were attenuated. This threshold dependent robustness supports a local Layer 4 circuit origin of the key domain specific dynamics and is now reported in the Results, Supplementary Figures, and Discussion. Second, we refined the interpretation of Layer 2/3 findings. While multi whisker versus single whisker differences in Layer 2/3 multi unit activity were reduced under more conservative spike detection criteria, single cell calcium imaging demonstrates that Layer 2/3 SST positive interneurons show stimulus dependent recruitment during multi whisker stimulation. We therefore reframed these data as evidence for cell type specific, stimulus dependent engagement of SST positive interneurons, without implying spatially focal barrel septa segregation at the level of Layer 2/3 population activity. Third, we further moderated mechanistic claims regarding Elfn1 knockout experiments. Throughout the revised manuscript, the SST Elfn1 mechanism is explicitly presented as a working model supported by converging anatomical, physiological, developmental, and genetic observations, rather than definitive causal proof. This framing is now emphasized in the Discussion. Finally, we expanded the Discussion to better contextualize our anatomical tracing results with prior work. We explicitly relate our retrograde labeling data to classic studies proposing parallel barrel and septa based processing streams, and extend these models by highlighting layer specific projection preferences in mouse whisker somatosensory cortex.

