## Supplementary material for "Local Inhibitory Dynamics Underpin Temporal Integration and Functional Segregation between Barrels and Septa in the Mouse Barrel Cortex": Response to Reviewers

Reviewer #1 (Public review):

Summary:
Argunşah et al. describe and investigate the mechanisms underlying the differential response dynamics of barrel vs septa domains in the whisker-related primary somatosensory cortex (S1). Upon repeated stimulation, the authors report that the response ratio between multi- and single-whisker stimulation increases in layer (L) 4 neurons of the septal domain, while remaining constant in barrel L4 neurons. The authors attribute this divergence to differences in short-term synaptic plasticity, particularly within somatostatin-expressing ($SST^{+}$) interneurons. This interpretation is supported by

1) the increased density of SST+ neurons in L4 of the septa compared to barrel domain,
2) the stronger response of (L2/3) SST+ neurons to repeated multi- vs single-whisker stimulation and
3) the reduced functional difference in single- versus multi-whisker response ratios across barrel and septal domains in Elfn1 KO mice, which lack a synaptic protein that confers characteristic short-term plasticity, notably in SST+ neurons.

Consistently, a decoder trained on WT data fails to generalize to Elfn1 KO responses. Finally, the authors report a relative enrichment of S2- and M1-projecting cell densities in L4 of the septal domain compared to the barrel domain, suggesting that septal and barrel circuits may differentially route information about single vs multi-whisker stimulation downstream of S1.

Strengths:
This paper describes and aims to study a circuit underlying differential response between barrel columns and septal domains of the primary somatosensory cortex. This work supports the view these two domains contribute distinctly to the processing single versus multi-whisker inputs and highlight the role of SST+ neuron and their short-term plasticity. Together, this study suggests that the barrel cortex multiplexes whisker-derived sensory information across its domains, enabling parallel processing within S1.

Weaknesses:
Although the divergence in responses to repeated single- versus multi-whisker stimulation between barrel and septal domains is consistent with a role for $SST^{+}$ neuron short-term plasticity, the evidence presented does not conclusively demonstrate that this mechanism is the critical driver of the difference. The lack of targeted recordings and manipulations limits the strength of this conclusion: $SST^{+}$ neuron activity is not measured in L4, nor is it assessed in a domain-specific manner. The Elfn1 knockout manipulation does not appear to selectively affect either stimulus condition, domain or interneuron subtype. Finally, all experiments were performed under anesthesia, which raises concerns about how well the reported dynamics generalize to awake cortical processing.

We thank the reviewer for their careful reading of the manuscript and their balanced assessment of both its strengths and limitations. We acknowledge the reviewer's concerns regarding the lack of direct, layer- and cell-type–specific recordings and manipulations of $SST^{+}$ interneurons, as well as the use of anesthesia. As noted in the Discussion, these factors limit the extent to which causal mechanisms can be established and the degree to which the reported dynamics can be generalized to awake cortical processing. For this reason, we intentionally frame the Elfn1–SST mechanism as a working model supported by converging anatomical, developmental, physiological, and genetic evidence, rather than as definitive proof. We believe this conceptual framing appropriately reflects the scope of the current data while highlighting clear directions for future work.

Reviewer #2 (Public review):

Summary:
Argunsah and colleagues demonstrate that SST expressing interneurons are concentrated in the mouse septa and differentially respond to repetitive multi-whisker inputs. Identifying how a specific neuronal phenotype impacts responses is an advance.

Strengths:
(1) Careful physiological and imaging studies.
(2) Novel result showing the role of SST+ neurons in shaping responses.
(3) Good use of a knockout animal to further the main hypothesis.
(4) Clear analytical techniques.

Comments on revisions:
The authors have effectively responded to my initial critiques - I have no further concerns.

We thank the reviewer for their positive evaluation of our work and for recognizing the novelty of the findings, the careful physiological and imaging approaches, the use of the Elfn1 knockout model, and the clarity of the analytical framework. We are pleased that the reviewer has no further concerns and appreciates the contribution of this study to understanding the role of $SST^+$ interneurons in shaping sensory processing in the barrel cortex.

---

Reviewer #3 (Public review):

Summary:
This study investigates the functional differences between barrel and septal columns in the mouse somatosensory cortex, focusing on how local inhibitory dynamics (particularly involving $SST^+$ interneurons) may mediate temporal integration of multi- whisker (MW) stimuli in septa. Using a combination of in vivo multi-unit recordings, calcium imaging, and anatomical tracing, the authors propose a model in which Elfn1-dependent synaptic facilitation onto $SST^+$ interneurons contributes to the distinct sensory responses to MW input in barrels and septa, enabling functional segregation between these domains.

Strengths:
The study presents a thought-provoking and useful conceptual model for understanding sensory processing in the somatosensory cortex. While barrel columns have been widely studied, septal regions remain relatively understudied in mice. If septa indeed act as selective integrators of distributed sensory input, this would suggest a novel computational role for cortical microcircuits beyond the classical view focused on barrels. Although still hypothetical, the proposed model in which $SST^+$ interneurons contribute to domain-specific sensory responses between barrel and septal domains is intriguing and opens new avenues for investigating inhibitory circuit mechanisms.

Weaknesses:
The primary limitation of this study lies in the spatial and cellular specificity of the recording techniques. The physiological data rely predominantly on unsorted multi-unit activity (MUA) recorded with low-channel-count silicon probes. Because MUA aggregates signals from multiple neurons over a radius of approximately 50-100 μm (often wider than the typical septal width in mice), this approach makes it difficult to confidently isolate activity originating strictly from within septal domains. The manuscript would benefit from additional analyses to validate the spatial specificity of these recordings, such as systematically varying spike detection thresholds to test the robustness of domain attribution, as suggested by the reviewer. Furthermore, although the authors now appropriately frame their findings in the Elfn1 knockout mice as indirect evidence, it is worth emphasizing that the study lacks direct in

vivo, cell-type-specific recordings and manipulations to more definitively test the proposed mechanism.

We thank the reviewer for their thorough and constructive evaluation of the manuscript and for highlighting both the conceptual strengths of the study and its technical limitations. We agree that the spatial and cellular specificity of unsorted multi-unit recordings imposes inherent constraints on the interpretation of domain-specific activity, particularly given the narrow width of septal compartments in mice. As now clarified in the manuscript, we do not claim absolute cellular specificity of “septal” recordings but rather interpret them as septal-enriched populations. To directly address this concern, we performed additional threshold-based analysis demonstrating that the key domain-specific effects persist selectively in Layer 4 under stricter spike-detection criteria, supporting a local circuit origin of the critical findings. Further, the more stringent detection criteria (Suppl Fig 3A) collapse the divergence seen in Layer2/3 (Suppl Fig 4C), suggesting that this divergence arises in Layer 4, where SST+ interneuron distributions diverge between barrel and septa.

We further agree that the Elfn1 knockout results provide indirect, rather than definitive, evidence for causal involvement of $SST^+$ interneurons and therefore intentionally frame the Elfn1–SST mechanism as a working model supported by converging anatomical, physiological, developmental, and genetic observations. We believe this explicitly moderated interpretation appropriately reflects the scope of the current data while establishing a clear conceptual framework and motivation for future studies employing cell-type-specific recordings and manipulations to directly test the proposed mechanism.

----------

Recommendations for the authors:

Reviewer #3 (Recommendations for the authors):

Major comments

(1) Interpretation of "septal" recordings: The authors claim that the activity recorded from electrodes placed in the septa can be confidently attributed to septal neurons. In my previous review, I raised a major concern that such "septal" recordings likely include spikes from adjacent barrels, given the broad spatial resolution of MUA and the narrowness of the septa in the mouse S1. In fact, the intermediate properties observed in septal recordings from wild-type mice could be explained by a mixture of activity from principal and neighboring barrels-an interpretation that contrasts with the authors' conclusion. Upon reviewing the probe model used (A8x8-Edge-5mm-100-200-177), I noticed a discrepancy between the manufacturer's design and the schematic provided in the manuscript. The electrodes are located near the right edge of the probe rather than the center, suggesting that neurons in adjacent barrels could easily be sampled. In my previous review, I therefore suggested alternative approaches, such as calcium imaging, to more convincingly support the authors' claims. However, the revised manuscript does not include new experiments or additional analyses addressing this issue. Instead, the authors argue that using a high spike detection threshold (SD > 7.5) ensures that recorded activity originates from septal neurons, even though this value does not appear particularly conservative, as it was merely adopted from a previous study without justification in the present context. While I agree that a higher threshold may reduce contamination from distant sources, it does not guarantee that only septal neurons contribute to the signal. By nature, MUA reflects activity from multiple neurons within a radius of at least 50-100 µm. To more rigorously support the claim of spatial specificity, I strongly encourage the authors to reanalyze their existing dataset by systematically varying the spike detection threshold and quantifying how the properties and selectivity of detected units change. If neurons closer to the electrode indeed exhibit distinct domain-specific properties, they should become more prominent as the threshold increases. Such an analysis would strengthen the authors' interpretation and improve the manuscript's impact, even in the absence of new experimental data. Alternatively,

the authors could revise their claims to acknowledge that the "septal" electrodes likely record from a population that includes septal neurons as well as neurons located at the periphery of principal and adjacent barrels.

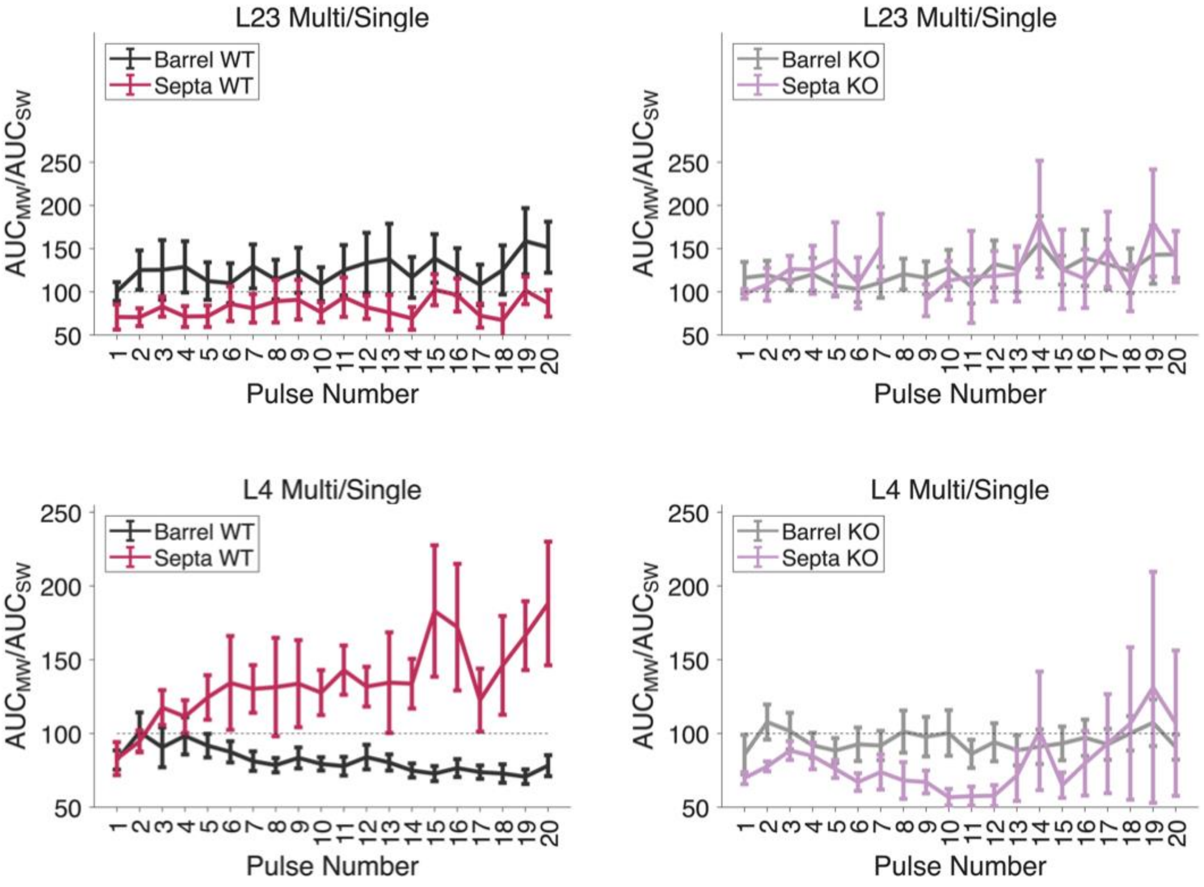


Figure 1. Spike detection with SD>9.5 and re-analysis of Layer2/3 and Layer 4 MW/SW data.

We agree with the reviewer that, by nature, MUA reflects the activity of multiple neurons within a spatial radius and that recordings obtained from electrodes positioned in the septa may include contributions from neurons located at the periphery of adjacent barrels. This concern is further compounded in superficial layers by probe geometry and orientation: given the narrow width of septa and the lateral spread of processes in upper cortical layers, recordings in L2/3 are inherently more susceptible to spatial mixing than those in layer 4, where columns are more compact and cytoarchitecturally distinct. To directly address these issues, we reanalyzed the same dataset using a more stringent spike detection threshold (SD > 9.5), compared to the originally reported SD > 7.5. Importantly, increasing the threshold selectively reduced or eliminated effects in L2/3, while the key domain-specific differences in L4 responses both the differential MW/SW dynamics in wild-type animals and their attenuation in Elfn1 knockout mice remained robust (Figure 1 above and the new Supp. Fig. 3. In the manuscript). This threshold-dependent dissociation is consistent with the interpretation that the critical effects reported in L4 arise from neurons spatially closer to the electrode and are less influenced by probe orientation or distant sources, rather than reflecting simple mixing of barrel signals. While this analysis does not claim absolute cellular exclusivity of septal neurons, it provides empirical support that the principal conclusions of the study are robust to stricter spatial sampling criteria and are particularly anchored in L4 circuitry. Accordingly, we now explicitly acknowledge in the manuscript that "septal" recordings likely represent septal-enriched populations rather than purely septal neurons, while emphasizing that the persistence of L4 effects under higher spike-detection thresholds strengthens the conclusion that local L4 inhibitory dynamics underlie the reported functional differences between barrel and septal domains.

The greater sensitivity of L2/3 results to spike-detection threshold is also expected based on both anatomical considerations and probe geometry. Neurons in L2/3 possess broader horizontal dendritic and axonal arbors and participate in more laterally distributed integration across columns, making population signals in these layers intrinsically less spatially focal. As a result, conservative spike-detection criteria preferentially suppress L2/3 effects, particularly when recordings are obtained with probes optimized for deeper layers. Importantly, our two-photon calcium imaging data while similarly limited to L2/3 demonstrate that $SST^+$ interneurons show locally measurable and stimulus-specific responses at the single-cell level, providing independent support that L2/3 $SST^+$ activity is stimulus-modulated rather than artifactual. Taken together, these observations suggest that L2/3 results reflect more distributed and integrative network activity, whereas the L4 effects that persist across thresholds are more directly attributable to local circuit mechanisms. This layer-specific dissociation further supports our interpretation that the central findings of the study are driven by local inhibitory dynamics in L4, with L2/3 activity reflecting downstream integration rather than primary domain-specific computation.

(2) Interpretation of the Elfn1 KO data: The authors' interpretation that Elfn1-dependent facilitation of $SST^+$ interneurons underlies the differential sensory responses between barrel and septal domains is conceptually appealing and supported by several converging, albeit indirect, lines of evidence. Specifically, the consistent correspondence among the differential activation of $SST^+$ neurons upon SWS and MWS, the late development of the barrel-septa differences in the responses to SWS and MWS, and the attenuation of this difference in Elfn1 knockout mice lends plausibility to the proposed model. However, it should be emphasized that the data remain indirect: the study does not include direct recordings of $SST^+$ neuronal activity from the knockout mice, nor cell-type- specific manipulations to demonstrate causal involvement. The mechanistic explanation therefore represents a hypothesis rather than definitive proof. That said, the authors clearly acknowledge these limitations in the Discussion and appropriately moderate their claims by presenting the SST-Elfn1 mechanism as a working model. Given this careful framing, the current manuscript can be regarded as a valuable conceptual contribution that advances our understanding of how inhibitory dynamics may shape temporal processing in the barrel cortex. Further experiments, as mentioned above, will be essential to test the causal role of this mechanism directly.

We thank the reviewer for this thoughtful and balanced assessment. We fully agree that the Elfn1 knockout experiments provide indirect rather than definitive evidence for a causal role of $SST^+$ interneurons in mediating the domain-specific MW/SW response dynamics between barrels and septa For this reason, throughout the revised manuscript we explicitly frame the Elfn1–SST mechanism as a working model rather than a proven mechanism.

Minor comments:

The authors have adequately addressed my previous minor comments. In this round, I carefully reviewed the revised manuscript and identified several issues related to references. I would also like to add a brief comment regarding the Discussion section:

(1) Stachniak et al., 2021 is included in the reference list but is not cited anywhere in the main text. Please either remove this entry or cite it appropriately in the manuscript.

Removed.

(2) Yamashita et al., 2018 is cited in the main text (Line 767), but it is not included in the reference list.

Fixed.

(3) Sylwestrak and Ghosh, 2012 is cited at Line 261 and Line 270, but likewise absent from the reference list.

Fixed.

(4) At Line 497, Chen et al., 2015 is cited, but, the appropriate and original reference would be Chen et al., 2013 (PMID: 23792559), which should either replace or precede the 2015 citation.

Added.

(5) At Line 221, El-Boustani et al., 2018 is cited. However, this study is based on the visual cortex, whereas the manuscript concerns the barrel cortex. A more relevant citation (e.g., Lefort et al., 2009 [PMID: 19186171]) would better support the discussion of cellular organization in the barrel cortex. Please consider updating the citation.

Thank you for this suggestion. We agree with the reviewer and now we have changed El-Boustani with Lefort et al. 2009 as suggested by the reviewer.

(6) Furthermore, Chakrabarti & Alloway (2006) performed tracer-based mapping of projections from barrel and septal columns in rat S1 and similarly suggested differential organization of M1- and S2-projection neurons in the barrel and septal regions.

Although the current study thoroughly analyzed the layer-specificity of the location of these projection neurons, the lack of explicit discussion of this relevant prior work is a notable omission.

The authors should incorporate a comparison with these results to better contextualize their findings.

The following text is added to the discussion: "Our retrograde labeling data supports and expands on previous work proposing similar models (Alloway, 2008; Chakrabarti and Alloway, 2006)."
